# Pre-existing cell subpopulations in primary prostate cancers display surface fingerprint of docetaxel-resistant cells

**DOI:** 10.1101/2021.01.28.428577

**Authors:** Stanislav Drápela, Barbora Kvokačková, Radek Fedr, Daniela Kurfürstová, Martin Morong, Vladimír Študent, Wytske M. van Weerden, Martin Puhr, Zoran Culig, Jan Bouchal, Karel Souček

## Abstract

**Background:** Docetaxel resistance represents a leading obstacle in the therapy of prostate cancer (PCa), resulting in lethal disease. Intratumoral heterogeneity, which is frequently driven by epithelial-mesenchymal plasticity significantly contributes to the limited treatment response, chemoresistance, and subsequent poor prognosis of patients with lethal PCa.

**Methods:** We employed a high-throughput flow cytometry screening to identify cell surface fingerprint that associates with docetaxel resistance in PCa cells. Using patient-derived xenografts, we validated protein expression of the most robustly changed antigens *in vivo* and further assessed this 6-molecule surface fingerprint in primary PCa tumors.

**Results:** We revealed the overexpression of SSEA-4 antigen in both *in vitro* and *in vivo* docetaxel-resistant models and confirmed the SSEA-4 enrichment in a subpopulation of freshly isolated primary PCa tumors. The level of ST3GAL2, an enzyme that is critically involved in the SSEA-4 synthesis, correlated with increased expression of CD44, CD59, and CD95 and reduced expression of EpCAM and CD9. SSEA-4 was further directly linked to the antimicrotubule agent resistance and poor prognosis in PCa patients.

**Conclusions:** We propose that the 6-molecule surface fingerprint associates with docetaxel resistance and pre-exists in a cell subpopulation of primary PCa tumors even before docetaxel treatment.

## Introduction

Docetaxel resistance represents a major obstacle in the therapy of prostate cancer (PCa) and frequently leads to disease relapse and death.^1^ Such resistance can generally arise from two different evolutionary pathways, either *de novo* from drug-tolerant “persister” cells during the therapy or as a consequence of pre-existing, resistant population driven by various mechanisms of intratumoral heterogeneity.^2,3^ The latter hypothesis is clinically considered as a *fait accompli* and posits that the rare resistant clones emerge in the tumor mass before treatment. These clones then drive the tumor relapse during treatment with targeted agents.^4,5^ The extent of such intratumoral heterogeneity is still poorly understood, and the clonality in the context of therapy resistance remains a critical bottleneck of current PCa research.^6–8^ Intratumoral heterogeneity is shaped to a significant extent by the lineage plasticity, which is under physiological conditions defined as the ability of a cell to reversibly or irreversibly modify its identity that differs in its original competence.^9,10^ In PCa cells, this context-dependent reprogramming of one committed phenotype is closely related to the formation of overt metastasis, acquisition of an invasive phenotype, and development of therapy resistance. This process is often accompanied by an epithelial-to-mesenchymal transition (EMT).^11–13^ Given the complex nature of these processes that potentially drive the development of docetaxel resistance, we hypothesize that the surfaceome of docetaxel-resistant cells shows distinct molecular features when compared to the cells that are therapy-sensitive.

We identified a 6-molecule surface fingerprint that reflects the phenotype of docetaxel-resistant PCa using *in vitro* models, exploiting an antibody-based, high-throughput platform for cell surface profiling. We validated the expression of these markers in docetaxel-resistant patient-derived xenografts (PDXs) *in vivo* and dissociated primary PCa tissues obtained after robotic prostatectomy. We revealed an upregulation of Stage-specific embryonic antigen-4 (SSEA-4), a glycosphingolipid, in both *in vitro* and *in vivo* docetaxel-resistant models and identified subpopulation cancer cells expressing high levels of SSEA-4 in fresh clinical specimens.

## Materials and Methods

### Cell lines, xenografts and chemicals

Docetaxel-resistant (DOC) DU145 and PC3 PCa cell lines were derived as previously reported.^14^ The docetaxel-sensitive PC346C and PC339 spheroid lines and their docetaxel-resistant derivatives, PC346C-DOC and PC339-DOC were established from PCa patient derived xenografts (PDXs) and propagated as described previously.^15,16^ All cell lines were maintained as defined in Supplementary Materials and Methods. The cells were routinely tested for mycoplasma contamination. The cells were authenticated using AmpFLSTR Identifiler Plus PCR Amplification Kit (TFS, Czech Republic) to verify their origin.

### Antibody-based cell surface screening and spectral flow cytometry

The detailed procedures of the antibody-based cell surface screening and spectral flow cytometry are described in Supplementary Materials and Methods. Briefly, for high-throughput cell surface screening cells were expanded, barcoded with CellTrace Violet or/and CellTrace DDAO (Far Red) amine-reactive fluorescent dyes, and subjected to the staining of 332 surface antigens using LEGENDScreen Human Cell Screening PE Kit (cat.no. 700001; Biolegend, San Diego, CA, USA). Data acquisition was performed on FACSVerse (Becton Dickinson (BD), Franklin Lakes, NJ, USA), accessed with Universal Loader. For multicolor spectral flow cytometry analysis, the single-cell suspensions from PDXs and patient samples were subjected to red blood cell lysis and stained using the cocktail of fluorochrome-conjugated primary antibodies. The selection of human cells in PDXs was done based on anti-human CD298 positivity.^17^ As for the patient samples, anti-human CD45, CD31, CD90 antibodies were used for the exclusion of leukocytes, endothelial-like cells and stromal cells.^18,19^ Samples were analyzed with SONY SP6800 Spectral Cell Analyzer (SONY, Japan). Acquired FCS files were exported and analyzed using FlowJo software (v10.0.7; BD). In all (spectral) flow cytometric experiments, dead cells were excluded from the analysis based on their positivity to LIVE/DEAD Fixable Dead Cell Stains (various dyes, Invitrogen, TFS). Cell aggregates and debris were excluded from the analysis based on a dual-parameter dot plot in which the pulse ratio (signal height/*y*-axis *vs.* signal area/*x*-axis) was displayed. The representative gating strategy is shown in Supplementary Figure S1A. Dilution, clonality, fluorochrome information, catalog numbers of the antibodies used for spectral flow cytometric analyses are provided in Supplementary Materials and Methods and Supplementary Table S4. All antibodies were titrated before use or used as recommended by the manufacturer.

### Patient-derived xenograft and prostate cancer tissue processing

PDX spheroids were harvested, dissociated and inoculated subcutaneously into the right flank of six-week-old male SHO mice. After 3 weeks, mice were euthanized and tumors were surgically excised, enzymatically dissociated using digestion media and proceeded for staining. All European Union Animal Welfare lines (EU Directive 2010/63/EU for animal experiments) were respected. A detailed description of *in vivo* xenografting and tumor dissociation is stated in Supplementary Materials and Methods. Animal experiments were approved by the Ethical Committee of IBP CAS and REKOZ, Academy of Sciences of the Czech Republic (AVCR 65/2016), supervised by the local ethical committee and performed by certified individuals (SD and KS). Fresh PCa tumor samples (50-150 mg), evaluated by licensed pathologists, were obtained from University Hospital Olomouc from patients undergoing robotic prostatectomy. All human tissue samples were obtained based on approval of the University Hospital Olomouc Ethical committee (Ref. No. 83/19) from donors that signed written informed consent. Tissue samples were minced, enzymatically digested and stained as described in Supplementary Materials and Methods.

### Data analysis

Regarding *Antibody-based cell surface screening* cell lines were deconvolved based on the signal of the fluorescent barcode as described before.^20^ Both medians of fluorescence and the percentage of positivity for PE channel were exported and analyzed. As for *Spectral flow cytometry analyses*, spectral overlaps were calculated and compensated based on spectral unmixing algorithms in SONY SP6800 Software (SONY, Japan). All data including multidimensional data from single-cell analyses and visualizations were analyzed in FlowJo (v10.0.7; BD). The multiparametric data were processed using multidimensional reduction algorithms including tSNE, UMAP, TriMap, FlowSOM, PhenoGraph and X-shift. Statistical analyses were performed in Prism (v5, GraphPad, La Jolla, CA, USA). Clinical data sets (accession numbers GSE21032 and TCGA-PRAD) were retrieved via https://portal.gdc.cancer.gov/projects/TCGA-PRAD, GEO (NCBI), Oncomine (TFS) and Genomic Data Commons Data Portal (NCI). Kaplan–Meier plots were assessed via customized analysis in DriverDBv3 cancer omics database^,21^ the median was set as a cutoff. Biocarta Pathway (Harmonizome) was used for the correlation analysis of z-score of sets of the proteins participating in the relevant pathway.^22^ Heat map generation was performed with Morpheus (Broad Institute, Cambridge, MA, USA).

## Results

### Docetaxel-resistant cell lines express distinct surface molecule profile

The surface characteristics of docetaxel-resistant PCa cells are, to date, unknown. We have adapted a high-throughput flow cytometry screening platform of cell surfaceome using a commercially available antibody screening platform in tandem with fluorescent barcoding of nine cell line models (Fig 1A and S1A).^20^ Further analysis of the surface signature of two well-described docetaxel-resistant cell lines DU145 DOC and PC3 DOC revealed that 81 of 332 antigens (~24%) were expressed on the surface of at least one cell model (Supplementary Figure S1B and Supplementary Tables S1 and S2). 14 of these antigens were strongly upregulated in both docetaxel-resistant cell lines, while only EpCAM was upregulated in both docetaxel-sensitive models (Figure 1B and 1C). Considering the list of surface molecules upregulated in the docetaxel-resistant cell lines, the 12 most robustly upregulated antigens - CD9, CD44, CD59, CD63, CD70, CD71, CD81, CD95, CD97, CD166, CD201 and SSEA-4 - were, along with EpCAM, selected for further profiling (Figure 1C and Supplementary Figure S1C). Subsequent data analysis showed, as expected, clustering of cells based on sensitivity to docetaxel, highlighting the gain of identical phenotypic features that can be attributed to the acquisition of docetaxel resistance (Figure 1D).

**Figure 1.**
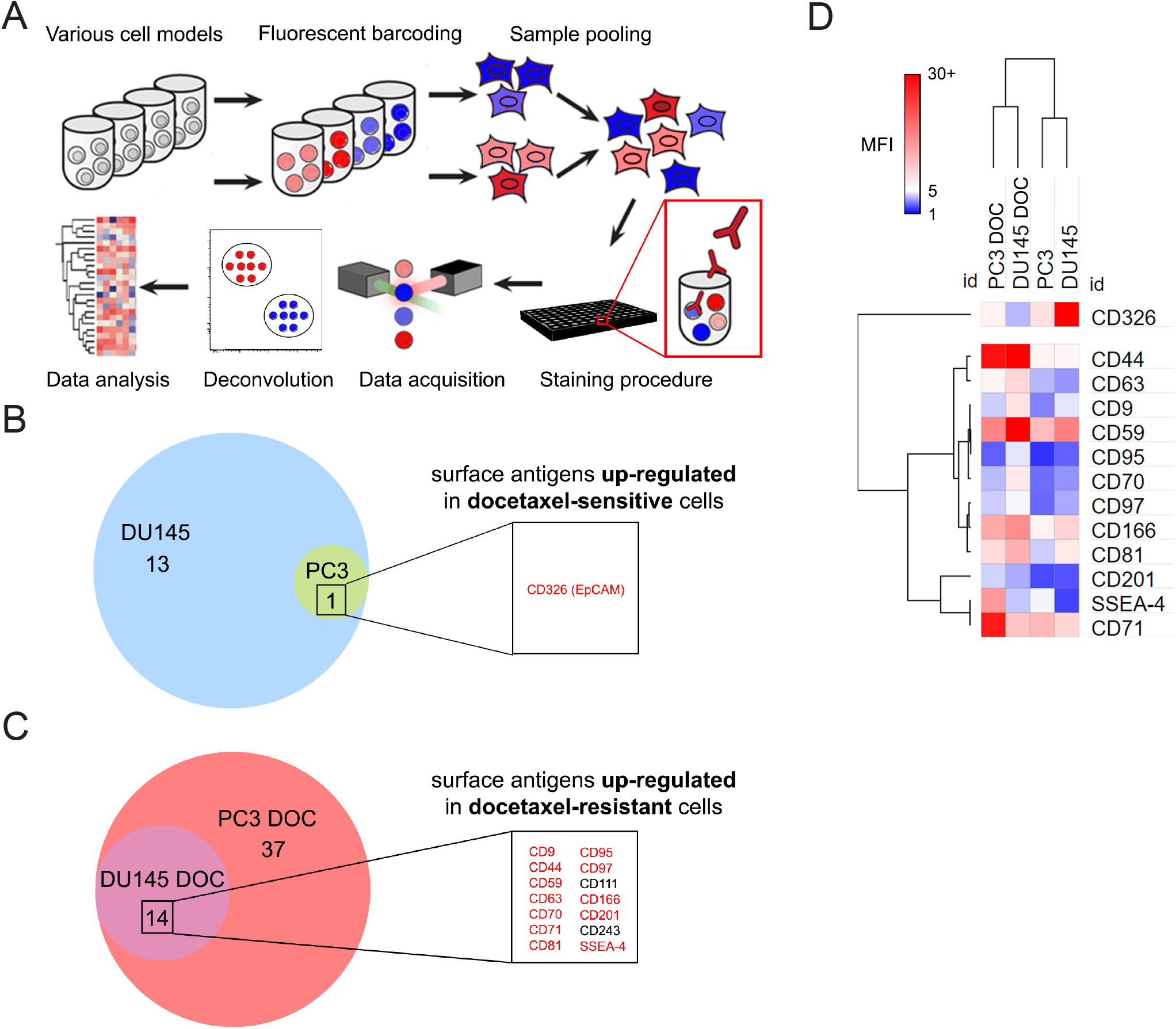
High-throughput profiling of surface molecules deregulated in docetaxel-sensitive and docetaxel-resistant PCa models *in vitro*. (A) The experimental workflow of fluorescent barcoding and high-throughput flow cytometry surface markers screening including data acquisition, deconvolution of cell lines based on the fluorescent barcode followed by data analysis. (B) Antigens upregulated on the surface of docetaxel-sensitive cells. (C) Antigens upregulated on the surface of docetaxel-resistant cells. Antigens selected for further validation are highlighted in red. The threshold for selection was set as >1.5-fold change. (D) Heatmap representing clustering (Spearman rank correlation) of cell lines and selected antigens based on the median fluorescence index (MFI). All figures represent data from screening analysis (n=1).

### *In vivo* validation of surface signature confirms a unique immunophenotype of docetaxel-resistant cells

We used docetaxel-resistant PC346C and PC339 preclinical PDX models^15,16^ to validate the results from *in vitro* screen using *in vivo* settings. The positive selection of human cells in the PDXs was confirmed by hCD298 staining. Multicolor spectral flow cytometry followed by the multidimensional data analysis revealed remarkable heterogeneity in the level of selected surface antigens in both models (Figure 2A). Except for CD71 (expressed at undetectable levels) and EpCAM, the expression of all remaining antigens-of-interest was significantly altered at least in one of the docetaxel-resistant PC346C DOC or PC339 DOC *ex vivo* xenografts, when compared to their docetaxel-sensitive counterparts (Figure 2B). Levels of surface CD9 were reduced in both docetaxel-resistant PDX models. In contrast, levels of CD44, CD59, CD63, CD81, CD97, CD201, and SSEA-4 antigens were upregulated in both docetaxel-resistant PDX models (Figure 2A and 2B). Due to the inconsistent trends in CD70 and CD166 expression between both PDX pairs, these molecules were excluded from further analysis (Figure 2B). Collectively, we determined a unique surface fingerprint that shapes the docetaxel-resistant phenotype in preclinical *in vivo* models of lethal PCa.

**Figure 2.**
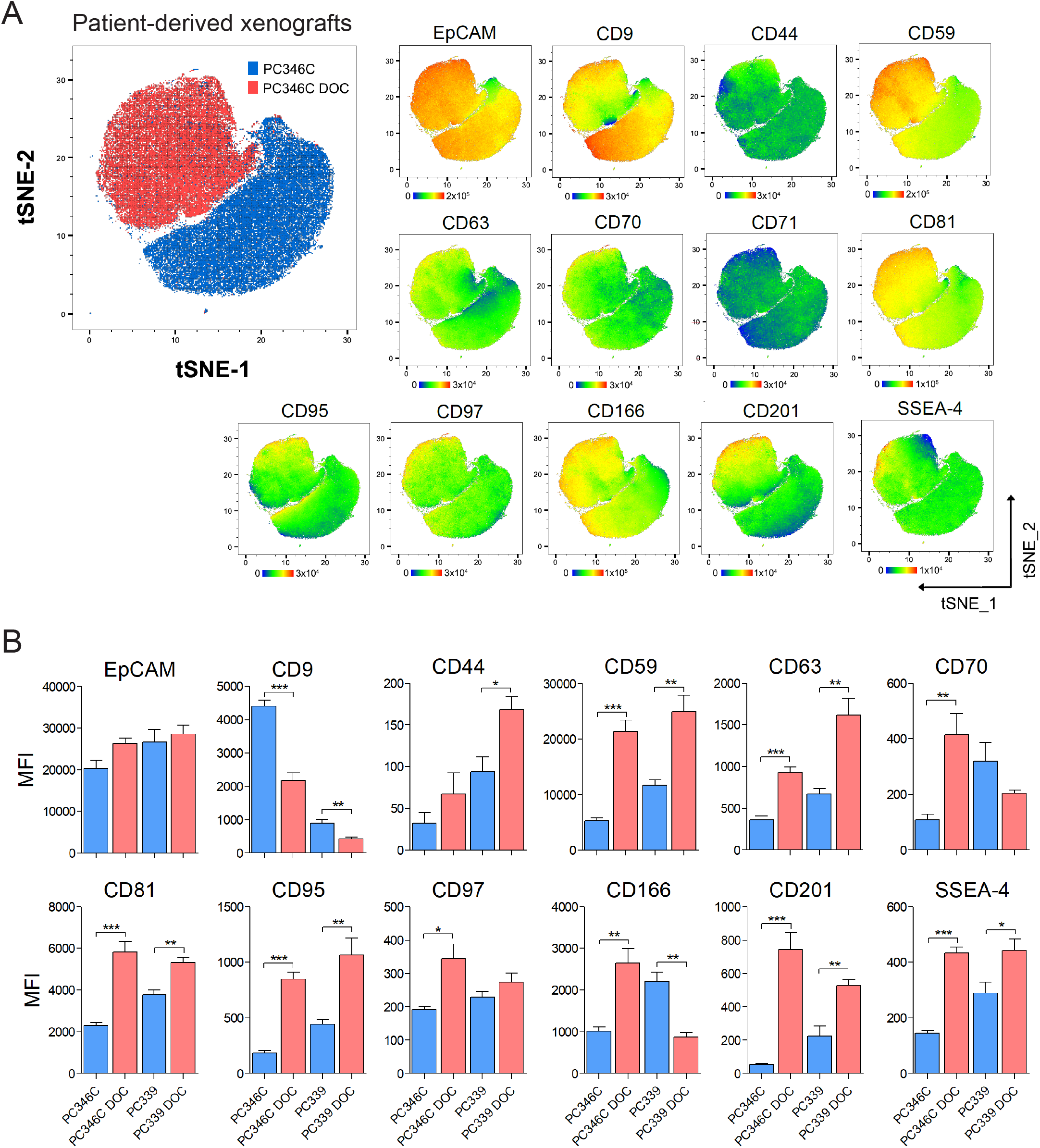
Validation of *in vitro* findings using PCa patient-derived xenografts. Tumors derived from PDXs were collected, dissociated and single-cell suspensions were stained as described in Materials and Methods. (A) The main tSNE plot (left) shows the distribution of docetaxel-sensitive PC346C (blue) and docetaxel-resistant PC346C-DOC (red) cells within the map. Related tSNE plots (right) illustrate the expression profile of particular antigens within the PC346C and PC346C-DOC clusters. The color corresponds to the fluorescence intensity (red - high; blue - low). The range of fluorescence intensity related to a particular antigen is depicted below each plot. (B) Expression profile of particular antigens in PDX models. Y-axes indicate the median fluorescence intensity. Data represent mean ± SEM from four independent tumors. ***, P<0.0001; **, P<0.01; *, P<0.05 by unpaired <*t*-test.

### A six-molecule fingerprint of docetaxel resistance identifies pre-existing cancer cell population in primary prostate clinical samples

We further profiled a cohort of eight untreated primary PCa patient samples using these nine antigens associated with docetaxel resistance *in vivo*, in combination with the epithelial marker EpCAM (Supplementary Table S3). The specimens were collected from radical prostatectomy and dissociated using a well-established protocol (*see* Supplementary Methods). Staining against CD45, CD31 and CD90 was used to exclude leukocytes, endothelial cells, and cancer-associated stromal cells. This approach allowed us to interrogate the expression of antigens-of-interest selectively on the surface of primary tumor epithelial cells. Advanced, multidimensional data analysis from patient samples revealed heterogeneity in the expression of all antigens, except for CD81 and CD97 which were not expressed in any of the patient samples and thus further considered as clinically irrelevant (Figure 3A). Unsupervised FlowSOM clustering algorithms uncovered a unique cell subpopulation (Pop3), characterized by the high expression levels of CD44, CD59, CD95, and SSEA-4 and low levels of EpCAM and CD9, mirroring the findings from docetaxel-resistant PDXs. Additionally, our analysis also revealed a cell subpopulation (Pop1) with a reversed phenotype that expressed low levels of CD44, CD59, CD95, and SSEA-4 and high levels of EpCAM and CD9, resembling the surfaceome of docetaxel-sensitive PDXs (Figure 3B and 3C and Supplementary Figure S2A-2C). Due to the very low to absent expression of CD63 and CD201 within these two patient subpopulations (Figure 3C), we omitted these antigens from our further investigations. Considering the previous results and expression profile of the 6-molecule surface fingerprint containing EpCAM, CD9, CD44, CD59, CD95 and SSEA-4, we evaluated PCa patient sample subpopulations Pop3 and Pop1 as potential prediction markers of sensitivity to docetaxel treatment in PCa patients (Figure 3D). The existence of Pop3 subpopulation was further confirmed, based on the 6-molecule surface fingerprint, using multiple clustering and projection algorithms that enabled multidimensional data reduction (Supplementary Figure S3). Finally, we showed that the Pop3 subpopulation that reflects the docetaxel-resistant phenotype was present in 5 of 8 patient samples as overt or rare subpopulation ranging from 1.3% (PCa3) to 55.7% (PCa6), illustrating patient-to-patient variability (Figure 3E and Supplementary Figure S2D). Conclusively, using preclinical models of docetaxel resistance and primary PCa patient samples, we propose the 6-molecule surface fingerprint composed of EpCAM, CD9, CD44, CD59, CD95 and SSEA-4 as a candidate set of surface antigens which may potentially predict response to docetaxel in PCa patients, prior therapy.

**Figure 3.**
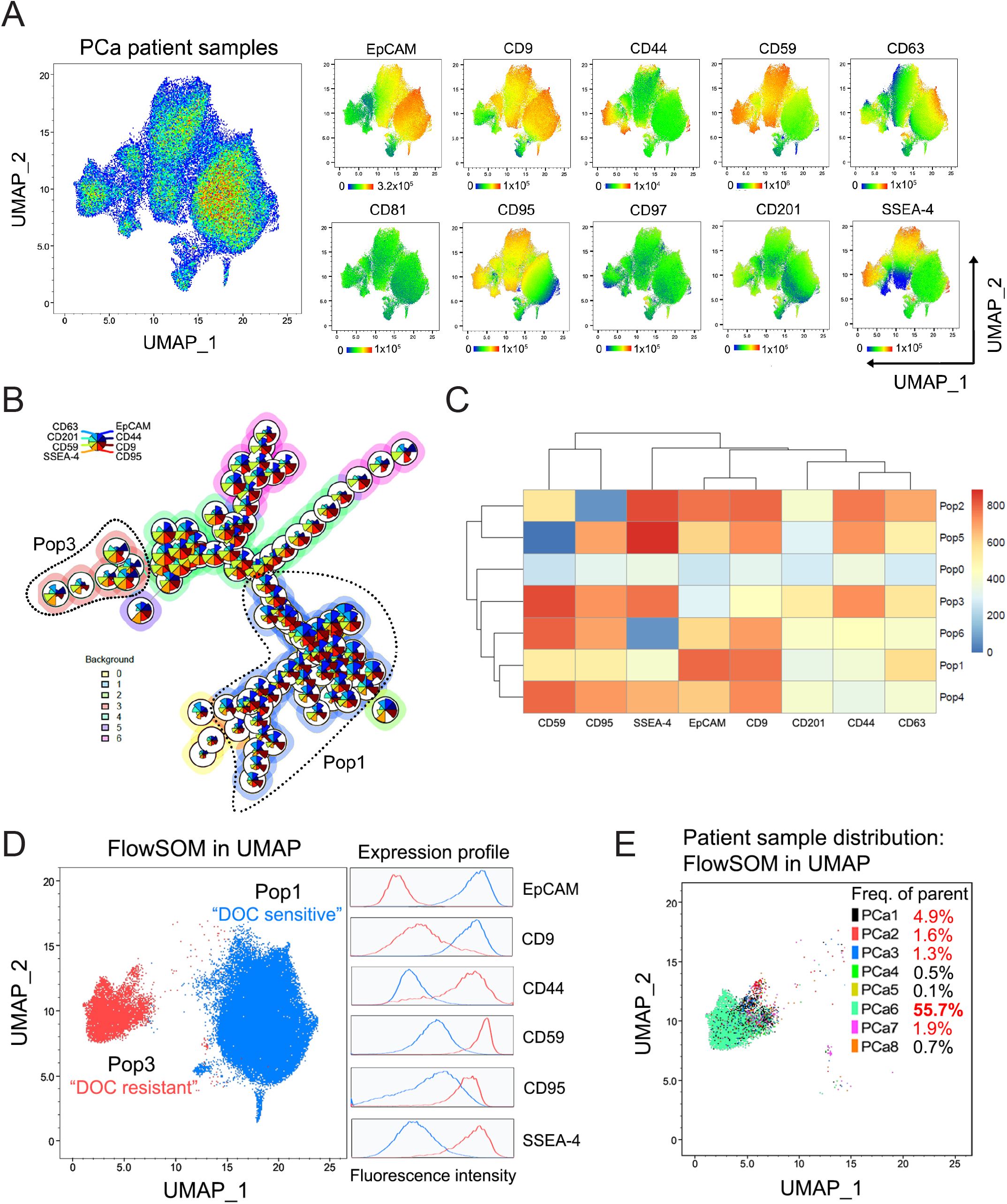
Inspection of the expression of selected antigens in dissociated primary prostate tumor samples. Fresh prostate cancer samples (n=8) collected from eight patients undergoing surgical prostatectomy were minced, enzymatically digested and stained as described in Supplementary Materials and Methods. (A) The main UMAP plot in pseudocolour (left) indicates the distribution of cells from eight prostate cancer patient samples within the map based on the expression of selected antigens. Related UMAP plots (right) illustrate the expression profile of particular antigens within the clusters. The color corresponds to the fluorescence intensity (red - high; blue - low). The range of fluorescence intensity related to a particular antigen is depicted below each plot. (B) FlowSOM algorithm illustrating clustering of cell subpopulation based on the eight markers with detectable expression in patient samples. FlowSOM is visualized as nodes, where each node represents a cluster of a specific subpopulation of cells, and pie-charts of the node reflect the contribution of different markers on to the phenotype of the cell cluster. (C) Heatmap corresponding to the FlowSOM clustering, depicting expression of particular antigens within each population. (D) UMAP visualization of selected Pop3 “DOC resistant” reflecting the fingerprint of docetaxel-resistant PDXs and Pop1 “DOC sensitive” reflecting the fingerprint of docetaxel-sensitive PDXs, determined by FlowSOM. Related expression profiles of the six most deregulated antigens are displayed by histograms on the right (red, “docetaxel-resistant”; blue, “docetaxel-sensitive”). (E) UMAP visualization of the distribution of patient samples within the “resistant” population of cells determined by FlowSOM. Each color denotes a different patient sample. The values in the legend refer to the % of cells belonging to the “resistant” population from the overall number of cells of each patient sample.

### Genes triggering SSEA-4 synthesis associate with poor survival and chemoresistance in prostate cancer patients

SSEA-4 was one of the most differentially expressed surface antigens discriminating docetaxel-resistant and docetaxel-sensitive cell lines and PDXs and further showed heterogeneous expression within primary PCa patient samples. We, therefore, hypothesized that its overexpression may play an important role in docetaxel resistance and subsequent poor disease outcome. We investigated the role of enzymes that are fundamental for SSEA-4 synthesis – ST3GAL2, ST3GAL1, and B3GALT5 (Figure 4A). Data from the TCGA dataset showed that increased expression of all three enzymes correlated with higher Gleason score and metastases (Supplementary Figure S4A-4C). High expression of ST3GAL2, ST3GAL1, and B3GALT5 indicated an unfavorable prognosis in the terms of survival probability, particularly in the progressive disease (Figure 4B and 4C). Importantly, ST3GAL2 expression positively correlated with the z-score of a geneset associated with resistance to antimicrotubule agents, particularly in patients with high-grade disease (GS≥8) (Figure 4D and 4E). These results thus demonstrate the clinical relevance of SSEA-4 in docetaxel-resistant PCa. Finally, our analysis of the TCGA dataset revealed a negative correlation between ST3GAL2 and EpCAM as well as CD9 expression, whereas CD44, CD59 and FAS (CD95) correlated positively (Figure 4F and Supplementary Figure S4D). Notably, a stronger correlation was observed in patients with high-grade disease (GS≥8), emphasizing the prominence of these markers in advanced PCa (Figure 4F, S4D). Collectively, these analyses endorse further clinical relevance of the 6-molecule surface fingerprint as predictive for docetaxel resistance in patients suffering from PCa.

**Figure 4.**
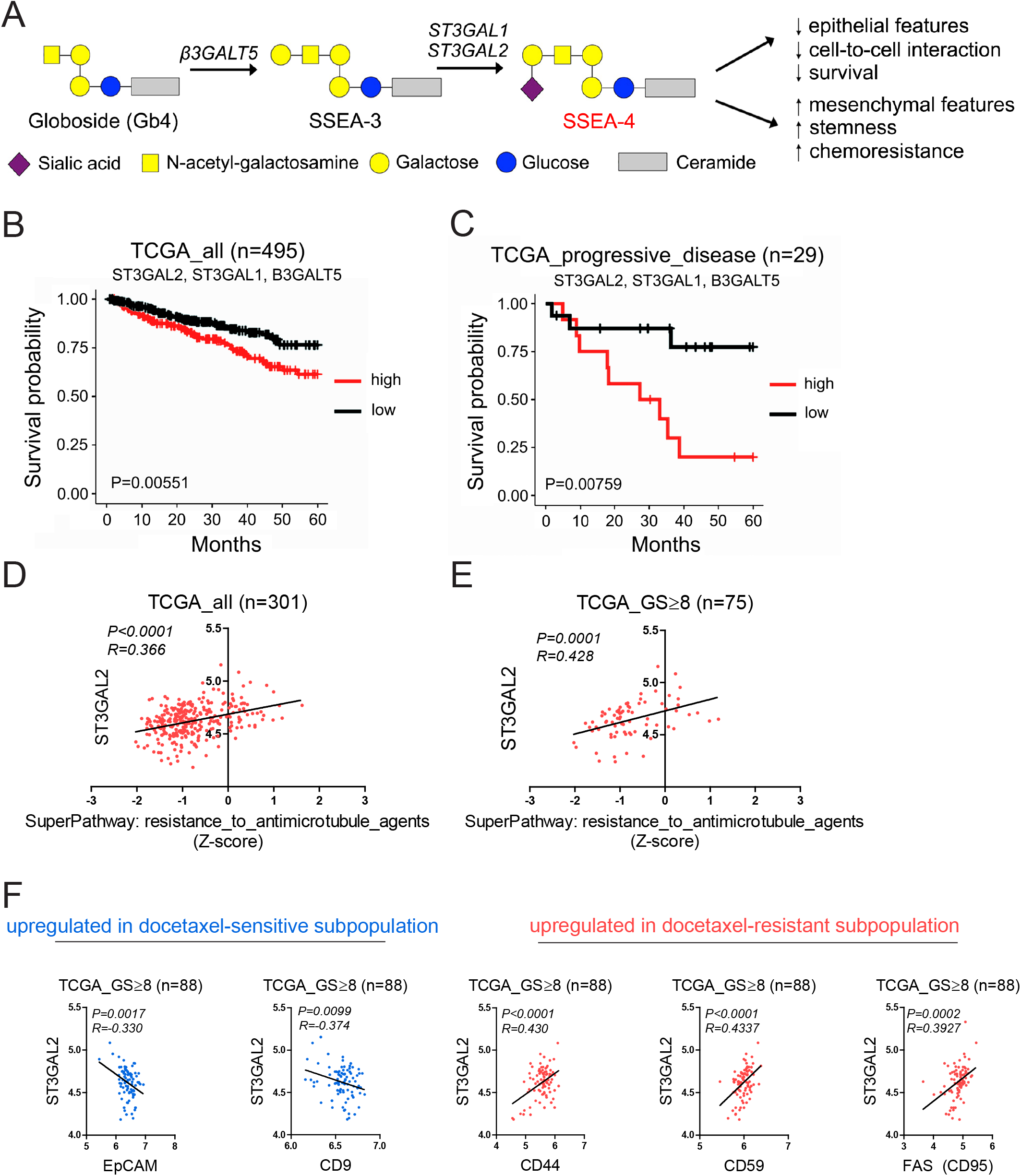
Correlation of SSEA-4 expression, resistance to microtubule agents and prognosis of high-grade PCa patients. (A) The scheme illustrates the precursors and enzymes implicated in the anabolism of SSEA-4 together with consequences of its overexpression. (B, C) The graphs show survival probability in a cohort of all (n=495) (B) or progressive disease only (n=29) (C) prostate cancer patients, respectively, according to high or low ST3GAL2, ST3GAL1 and B3GALT5 expression levels (TCGA). (D, E) Correlation analysis between ST3GAL2 and z-score of the genes implicated in the resistance to antimicrotubule agents (Biocarta Pathway-Harmonizome) using all available data from TCGA samples (n=301) (D) and data from high-grade (GS≥8) prostate cancer patients (n=75) (E). (F) Correlation analysis between ST3GAL2 and surface antigens in primary PCa patient samples, upregulated in likely DOC sensitive subpopulation (Pop1; in blue) or upregulated in likely DOC resistant subpopulation, (Pop3; in red), according to the PDX fingerprints. Data portray the expression of antigens in patients suffering from high-grade (GS≥8) prostate cancer disease (n=88) (TCGA).

## Discussion

Despite the initial therapy success, PCa progresses to the life-threatening stage of metastatic castration-resistant prostate cancer (CRPC) that remains the second major cause of cancer death in men.^23,24^ Docetaxel represents the backbone treatment against CRPC.^25–27^ However, its clinical utility is compromised in a significant proportion of men due to the development of acquired therapy resistance.^28^ Recent therapeutic approaches utilize early-stage administration of docetaxel alone as well as in combination with hormonal therapy or androgen-deprivation in hormone-sensitive PCa patients.^29,30^ Despite that the docetaxel-resistance represents such a critical obstacle in PCa therapy with lethal consequences, specific changes in the transcriptome and proteome of docetaxel-resistant cells remain largely uncovered. Here, we investigated the tumor cell phenotype linked to the docetaxel resistance using the analysis of surfaceome signature *in vitro* and *in vivo*, followed by validation in primary PCa patient samples and publicly available clinical datasets.

We used well-described docetaxel-resistant *in vitro* models DU145 DOC and PC3 DOC for the identification of the initial *in vitro* surface signature.^14^ These cells were derived by exposure of parental cells to the increasing concentrations of docetaxel until resistance occurred.^14^ Previous studies using these cell lines suggested EMT, including upregulation of vimentin, CD44, or ZEB1 and loss of E-cadherin, as a potential driving mechanism of docetaxel resistance.^14,31,32^ Additionally, CD44 was also established as a driver of invasion and migration and its overexpression was associated with therapy resistance and neuroendocrine-like phenotype in PCa.^33,34^ In concordance with these reports, our screen revealed downregulation of the epithelial marker EpCAM and upregulation of the mesenchymal marker CD44 in docetaxel-resistant models, suggesting an EMT switch. Besides, 11 antigens were upregulated in both DU145 DOC and PC3 DOC cells. *In vivo* validation of these antigens using docetaxel-resistant PC346C-DOC and PC339-DOC PDX models^15,16^ revealed significant deregulation in 9 of them. A consistent pattern of up- or downregulation was observed in CD9, CD44, CD59, CD63, CD81, CD95, CD97, CD201 and SSEA-4 antigens in both docetaxel-resistant PDXs as compared to their docetaxel-sensitive counterparts. The role of tetraspanin family members CD9, CD63 and CD81 was previously discussed in the context of tumor stage, chemoresistance and patient outcome in several studies.^35^ CD9 is a tumor suppressor associated with better prognosis^20^ that has also manifested oncogenic and pro-metastatic functions eventually resulting in chemoresistance.^36,37^ Overexpression of CD63 and CD81 was associated with EMT, chemoresistance and poor prognosis in various types of malignancies, including PCa.^38–40^ CD59 is a complement regulatory protein that was shown to be overexpressed in 36% of prostate adenocarcinomas and corresponded to the earlier biochemical relapse and adverse patient prognosis.^41^ CD95 (APO-1/Fas), a membrane protein belonging to the TNF receptor superfamily, contributes to the tumor-promoting effects accompanied by EMT and metastatic phenotype in oxaliplatin-resistant cells^42^ and its expression in primary PCa cells was, similarly to study, strongly stimulated by docetaxel treatment.^43^ The upregulation in CD97 and CD201 antigens has never been linked to chemoresistance before but was reported to correlate with PCa invasiveness.^44,45^ Due to the lack of mechanistic studies, the essential role of these surface molecules in docetaxel-resistant PCa remains controversial.

SSEA-4, which is also known as MSGG or LKE antigen, is a globo-series ganglioside that was identified as a promising biomarker and potentially a druggable target for the therapy of docetaxel-resistant PCa. Here, we show that SSEA-4 is uniformly upregulated in all *in vitro* and *in vivo* models of docetaxel resistance and is overexpressed in a subpopulation of primary tumor epithelial cells. Notably, SSEA-4 expression correlated with resistance to antimicrotubule agents, high-grade disease and unfavorable prognosis in PCa patients. This result goes hand-in-hand with the previously published studies associating SSEA-4 with tumorigenicity, chemoresistance and mesenchymal features in various malignancies.^46–48^ Aloia et al. have suggested that SSEA-4 might be induced by chemotherapy in a subpopulation of tumor cells.^47^ Our data also demonstrate that SSEA-4 correlates with the EMT status of PCa cells. This phenomenon was previously described in several studies, where SSEA-4 expression defined spontaneous loss of epithelial phenotype, characterized by the loss of cell-to-cell interactions and gain of mesenchymal features.^20,49^ Additionally, concurrent overexpression of SSEA-4 and CD44 in a subpopulation of tumor cells displayed extensive cancer stem-like cell characteristics, malignant behavior and worse overall survival.^50–52^ Given the current knowledge, such cells seem to be predisposed for chemoresistance and can be responsible for the induction of intratumoral heterogeneity and metastatic spread.^53,54^ Importantly, due to its rare expression in pluripotent human embryonic stem cells and limited expression in normal tissue, SSEA-4 was also proposed as an attractive target for CAR T cell and other immune-based therapies.^48,52,55^

Validation of the surface fingerprint, defined in preclinical PDX models, in a cohort of PCa patient samples resulted in a 6-molecule surface fingerprint composed of EpCAM, CD9, CD44, CD59, CD95 and SSEA-4. This fingerprint led to the identification of the Pop3 subpopulation, harboring key docetaxel resistance markers identified *in vivo*, and may indicate docetaxel-resistant tumor subpopulation in patients. The functional role and impact of these molecules on the docetaxel resistance and sensitivity have to be further addressed. This 6-molecule surface fingerprint expression profile revealed a remarkably, but not surprisingly high patient-to-patient variability. Importantly, the distribution and abundance of the cell subpopulation with docetaxel-resistant signature were not dependent on the disease stage and previous therapy history (Supplementary Table S3). This finding suggests that docetaxel resistance is shaped by a pre-existing intratumoral heterogeneity, as a consequence of early tumor plasticity, rather than *de novo* risen from drug-tolerant “persister” cells.

Despite that the function and contribution to docetaxel resistance of these six molecules remain yet to be uncovered, we pose that this unique fingerprint can serve in the clinical settings for early identification of docetaxel-resistant cells in PCa patients. Further, detailed classification of the subpopulations identified in our study using advanced techniques, such as single-cell transcriptomics and proteomics, will reveal the mechanisms responsible for docetaxel resistance and ultimately lead to the improved clinical management of aggressive prostate cancer.

## Abbreviations

*CAR*: chimeric antigen receptor
*CRPC*: castration-resistant prostate cancer
*DOC*: docetaxel-resistant
*EMT*: epithelial-to-mesenchymal transition
*PCa*: prostate cancer
*PDXs*: patient-derived xenografts
*SSEA-4*: stage-specific embryonic antigen-4

## Authors’ Contributions

SD performed experiments, analyzed and interpreted the data, and wrote and reviewed the manuscript. BK assisted with the data acquisition and interpretation. RF provided technical support. DK and MM performed a pathological evaluation of the normal prostate and prostate cancer tissue. VŠ performed surgical interventions and tumor excision. JB wrote and reviewed the manuscript. KS conceptualized and designed the study, interpreted the data, reviewed the manuscript, and supervised the study. WMvW, MP, and ZC provided the experimental models and reviewed the manuscript. All authors read and approved the final version of this manuscript.

## Acknowledgement

The authors would like to thank Iva Lišková, Martina Urbánková and Kateřina Svobodová for technical assistance; Drs. Nina Tokanová and Ráchel Víchová for the maintenance of the animal facility; Dr. Gvantsa Kharaishvili and Mgr. Monika Levková for help with the collection of patient samples; Drs. Ján Remšík and Jiřina Procházková for critical reading and thoughtful discussions and Dr. Eva Slabáková for proofreading the manuscript. The results published here are in part based upon data generated by the TCGA Research Network: https://www.cancer.gov/tcga.

## Ethical approval and consent to participate

All human tissue samples were obtained at University Hospital Olomouc, Olomouc, Czech Republic based on approval of the University Hospital Ethical committee (Ref. No. 83/19) from donors that signed written informed consent.

## Availability of data and materials

Preprocessed data from the high-throughput screening are available in Supplementary Information. Additional data and materials used in this study that are not available commercially are available upon reasonable request from the lead contact. Human tissues are unique biological resources that are not available for further distribution. Further information and requests for resources and reagents should be directed to and will be fulfilled by the Lead Contact, Karel Souček.

## Supplementary Information

### Supplementary Material and Methods

#### Cell lines, xenografts and chemicals

Docetaxel-resistant (DOC) DU145 and PC3 prostate cancer cell lines were derived as previously reported.^1^ Docetaxel resistance was maintained by continuous supply of docetaxel (cat.no. 9886, Cell Signaling, Danvers, MA, USA) in the final concentration of 12.5 nM. Cell lines were maintained at 37°C (5% CO2) in RPMI 1640 media (cat.no. 72400-021, Thermo Fisher Scientific (TFS), Waltham, MA, USA) supplemented with 10% fetal bovine serum (cat.no. 10270106, TFS, USA) and 100 U/mL penicillin/streptomycin (cat.no. XC-A4122/100, Biosera, France). The PDXs were established and cultured as described previously.^2,3^ Docetaxel resistance was maintained by continuous supply of docetaxel in the final concentration of 1 nM. PDXs were maintained at 37°C (5% CO2) in DMEM/F12 1:1 media (cat.no. 31330, TFS, USA) supplemented with 2 % fetal bovine serum and 100 U/mL penicillin/streptomycin, 0.01% BSA (cat.no. 11930, Serva, Germany), 10 ng/mL rhEGF (cat.no. E9644, Merck, Germany), 1 % ITS-G (cat.no. 41400-045, TFS, USA), 0.5 ug/mL hydrocortisone (cat.no. H0888, Merck, Germany), 1 nM triiodothyronine (cat.no. IRMM469, Merck, Germany), 0.1 mM O-phosphorylethanolamine (cat.no. P0503, Merck, Germany), 50 ng/mL cholera toxin (cat.no. C8052, Merck, Germany), 0.1 μg/mL fibronectin from human plasma (cat.no. F0895, Merck, Germany), 20 μg/mL fetuin from fetal bovine serum (cat.no. F3385, Merck, Germany) and 0.1 nM methyltrienolone (R1881) (cat.no. 816439, MolPort, Latvia).

#### Antibody-based cell surface screening

For surface profiling, cell lines were expanded in appropriate medium and harvested by incubation in 1.35 mM EDTA solution in PBS to allow non-enzymatic weakening of the cell junctions (2 mins for DU145 and DU145 DOC and 3 mins for PC3 and PC3 DOC cells) followed by mild HyClone HyQTase (cat.no. SV3003001, GE Healthcare Bio-Sciences AB, Sweden) incubation for 5 mins, as some antigens included in the screen are sensitive to massive trypsinization and differences in cell harvesting would obscure the results. HyClone HyQTase was then neutralized with cell culture medium containing 10% FBS (TFS, USA). Suspension of each cell line was counted, washed with PBS and 5×10^7^ cells were barcoded with CellTrace Violet or/and CellTrace DDAO (Far Red, TFS, USA) amine-reactive fluorescent dyes diluted in PBS (1 mL staining solution per 1×10^7^ cells), for 15 min at 37°C while gently shaking on the vertical rotator. Remaining unreacted dye was then quenched by addition of a cell culture medium containing 10% FBS and incubation for 30 min at 37°C in the dark. Cell suspensions were then washed with PBS and stained with LIVE/DEAD Green Fixable Dead Cell Stain diluted 1:1000 in PBS for 15 min at 4°C (1 mL staining solution per 1×10^7^ cells; TFS, USA). The cells were washed with PBS, filtered via 100 μm cell strainer to remove large cell aggregates, re-counted and 45×10^7^ cells from each cell line were then pooled in 27 mL of Cell Stain buffer (part of LEGENDScreen Kit, cat.no. 700001; Biolegend, San Diego, CA, USA) and filtered via 70 μm cell strainer. Cell suspension (75 μL/well, equal to a pool of 0.75×10^6^ cells/well) was then dispensed into reconstituted LEGENDScreen Human Cell Screening PE Kit 96 well plates and incubated 20 min at 4°C in dark. After reconstitution with MQ water, each well contained 25 μL of single, validated and pre-titrated antibody conjugated with Phycoerythrin (PE). Following the staining, the plates were spun down 500g for 6 min and the supernatant was dumped by quickly inverting and flicking the plate. The cells were washed with a Cell Stain Buffer and fixed in a Fixation Buffer (part of LEGENDScreen Kit) for 10 min at RT as recommended by the manufacturer. The cells were washed twice, resuspended in Cell Stain Buffer and processed for analysis.

#### Xenograft mouse experiments and patient-derived xenograft tumor processing

Floating spheroids of PDXs cell cultures were harvested, washed with 1.35 mM EDTA solution in PBS and subjected to non-enzymatic weakening of the cell junctions using HyClone HyQTase for 10 min at 37°C. HyClone HyQTase was then neutralized with cell culture medium containing 10% FBS. The suspension of each cell model was filtered via 70 μm cell strainer to remove large cell aggregates, counted using CASY cell counter (OLS OMNI Life Science, Germany) and a total of 1×10^6^ of PDX cells were re-suspended in the 1:1 mix of ice-cold PBS and Matrigel (Corning Incorporated, NY, USA) and inoculated subcutaneously into the right flank (dorsally) of six-week-old male SHO mice. The experimental unit refers to single animal. Severe combined immuno-deficient mice (SCID) hairless outbred (SHO) (Crl:SHO-PrkdcscidHrhr) were obtained from Charles River Laboratories (Wilmington, MA, USA). Mice were housed in sterile cages in a temperature-controlled room with 12-h light–dark cycle. Each cohort (animals inoculated by particular PDX model) contained 4 mice, corresponding to 16 mice in total. No criteria for including and excluding animals during the experiments were applied. The confounders were not controlled. Mice were euthanized with CO_2_ 3 weeks after inoculation when 1cm^3^ big, determined by caliper measurement. The tumors were surgically excised and enzymatically dissociated using digestion media containing 2 mg/mL Collagenase Type I (cat.no. LS 004194, Worthington, Lakewood, NJ, USA) and 0.6 U/mL Dispase II (Roche, Switzerland) in RPMI 1640 for 1 hour at 37°C on a horizontal rotator. Samples were then treated with 15 μg/mL DNase I (Roche, Switzerland) for 5 min at 37°C, washed with sterile PBS, filtered through 100 μm and 50 μm cell strainers to remove large cell aggregates. After washing, red blood cells were lysed with ACK buffer (155 mM ammonium chloride, 10 mM potassium bicarbonate and 100 μM EDTA solution in sterile MQ water). Cells were counted using CASY cell counter and 1,5×10^6^ of cells were used for the staining of viability and surface marker expression. The animal protocol was prepared and approved before the experiment was started. All European Union Animal Welfare lines (EU Directive 2010/63/EU for animal experiments) were respected. Animal experiments were approved by the Ethical Committee of IBP CAS and REKOZ, Academy of Sciences of the Czech Republic (AVCR 65/2016), supervised by the local ethical committee and performed by certified individuals (SD and KS).

#### Prostate cancer tissue processing

Tissue samples were minced to 1 mm pieces and enzymatically digested in HBSS buffer (Hank’s Balanced Salt Solution + 0,035% NaHCO_3_) containing 2.5 mg/mL Collagenase type I, and 10 μM Y-27632 dihydrochloride (cat.no. sc-281642A, Santa Cruz Biotechnology, Dallas, TX, USA), for 2-4 hours (depending on tissue toughness) at 37°C. Gentle agitation (40 RPM) was used to prevent viable cell loss and non-specific surface epitope cleavage. Samples were then treated with 15 μg/mL DNase I for 5 min at 37°C, washed with sterile PBS, filtered through 50 μm cell strainer and subjected to staining in fresh condition.

#### Spectral flow cytometry analysis

For multicolour panel testing, the single-cell suspensions from PDXs and patient samples were washed with PBS and subjected to red blood cells lysis with ACK buffer followed by filtration via 50 μm strainer. All suspensions were further stained under non-sterile conditions with LIVE/DEAD Yellow Fixable Dead Cell Stain diluted 1:500 in PBS for 15 min at RT (100 μL staining solution per 1×10^6^ cells; TFS, USA) to exclude dead cells from the analysis. Next for PDXs, a cocktail of fluorochrome-conjugated primary antibodies containing anti-human CD298, CD9, CD44, CD59, CD63, CD70, CD71, CD81, CD95, CD97, CD166 (ALCAM), CD201, SSEA-4, EpCAM was used for extracellular staining in a single tube. For the patient samples, the same panel of antibodies (excluding CD298) was enriched by FITC-conjugated anti-human CD45, CD31, CD90 antibodies. Suspensions were stained in sodium azide (NaN3) buffer (0.02% sodium azide + 1% BSA in PBS) enriched by Super Bright Complete Staining Buffer (cat.no. SB-4401-42, TFS, USA) for 20 min at 4°C, then washed and processed to analysis.

#### Data analysis

As for *Antibody-based cell surface screening*, compensations were calculated automatically in BD FACSuite™ Software (BD) from single-conjugate stained UltraComp eBeads (eBioscience) and applied during data acquisition and analysis. Gates for positivity and isotype median of fluorescence were set based on isotype controls. For median fluorescence index (MFI) calculation, mean from all isotype controls for each cell line was used. Table showing the percentage of positivity and MFI for each cell line and each analyzed antigen is available as Supplementary Table S1 and S2. As for *Spectral flow cytometry analyses*, spectral overlaps were calculated and compensated based on the fluorochrome spectra uploaded into the Spectral Library using positive controls (cell lines) and following automatic spectral unmixing algorithm in SONY SP6800 Software (SONY, Japan). Fluorescence minus one (FMO) controls were measured for all fluorochromes and used for regular gating. For the data visualization using various algorithms listed below, the number of events was prior downsampled to 1×10^4^ per each patient sample. Such samples were concatenated and subjected to the multidimensional reduction algorithms. We used tSNE algorithm^4^ for mapping of high-dimensional cytometry data onto two dimensions. tSNE plotted individual cells in a visual similar to a scatter plot while using all pairwise distances in high dimension to determine each cell’s location in the plot. The tSNE plots are visualized in pseudocolour and each dot represents a single cell – the lowest expression of the selected marker is in dark blue and the highest expression in dark red, the corresponding scale is next to each plot and reflects the level of marker expression. Similar to tSNE, the Uniform Manifold Approximation and Projection (UMAP) algorithm, enabling nonlinear dimension reduction, was used to uniformly distribute data on Riemannian manifold.^5^ To further visualize multidimensional data in self-organizing maps, we applied the two-level clustering algorithm, FlowSOM.^6^ Briefly, in such visualization, each node represents a cluster of a specific subpopulation of cells and pie-charts of the node reflect the contribution of different markers to the phenotype of the cell cluster. Since FlowSOM enables an adjustable number of the clusters for visualization, K-finder algorithm was applied on selected parameters prior FlowSOM analysis to set “K” value and thus approximate the number of clusters in an unbiased way. The dimensionality reduction algorithm TriMap (not published, arXiv:1910.00204), and other clustering algorithms PhenoGraph^7^ and X-shift^8^, together with ClusterExplorer were used to validate the presence of FlowSOM-identified clusters. Only viable, single cells were included in analyses.

## Supplementary Figures

**Figure S1.**
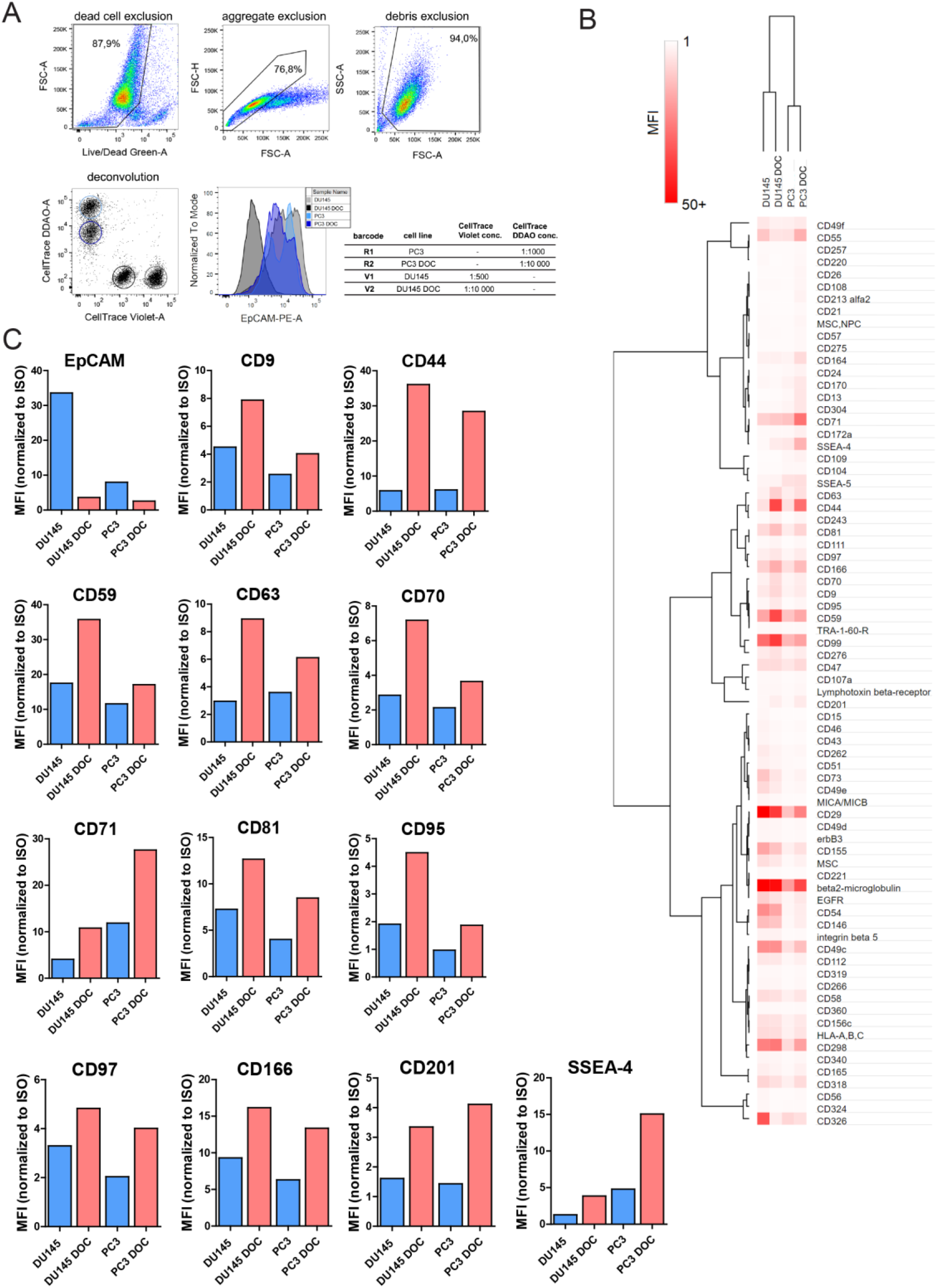
High-throughput profiling of surface molecules deregulated in docetaxel-resistant PCa models *in vitro*. (A) The scheme shows a full gating strategy (removal of dead cells, cell aggregates and debris) used for the analysis of all flow cytometric data in this manuscript and the deconvolution of fluorescent barcoding, used for the high-throughput screen. Detailed information is provided in Supplementary Materials and Methods. (B) A heatmap shows median fluorescence index (MFI) for all models and all antigens that were identified on the surface of at least one cell line. (C) Expression profile of antigens that were robustly changed in both DU145 and PC3 docetaxel-resistant models compared to their docetaxel-sensitive counterparts. Y-axis indicates the median fluorescence index (median fluorescence intensity of the specific antigen normalized to the median fluorescence intensity of ISO control).

**Figure S2.**
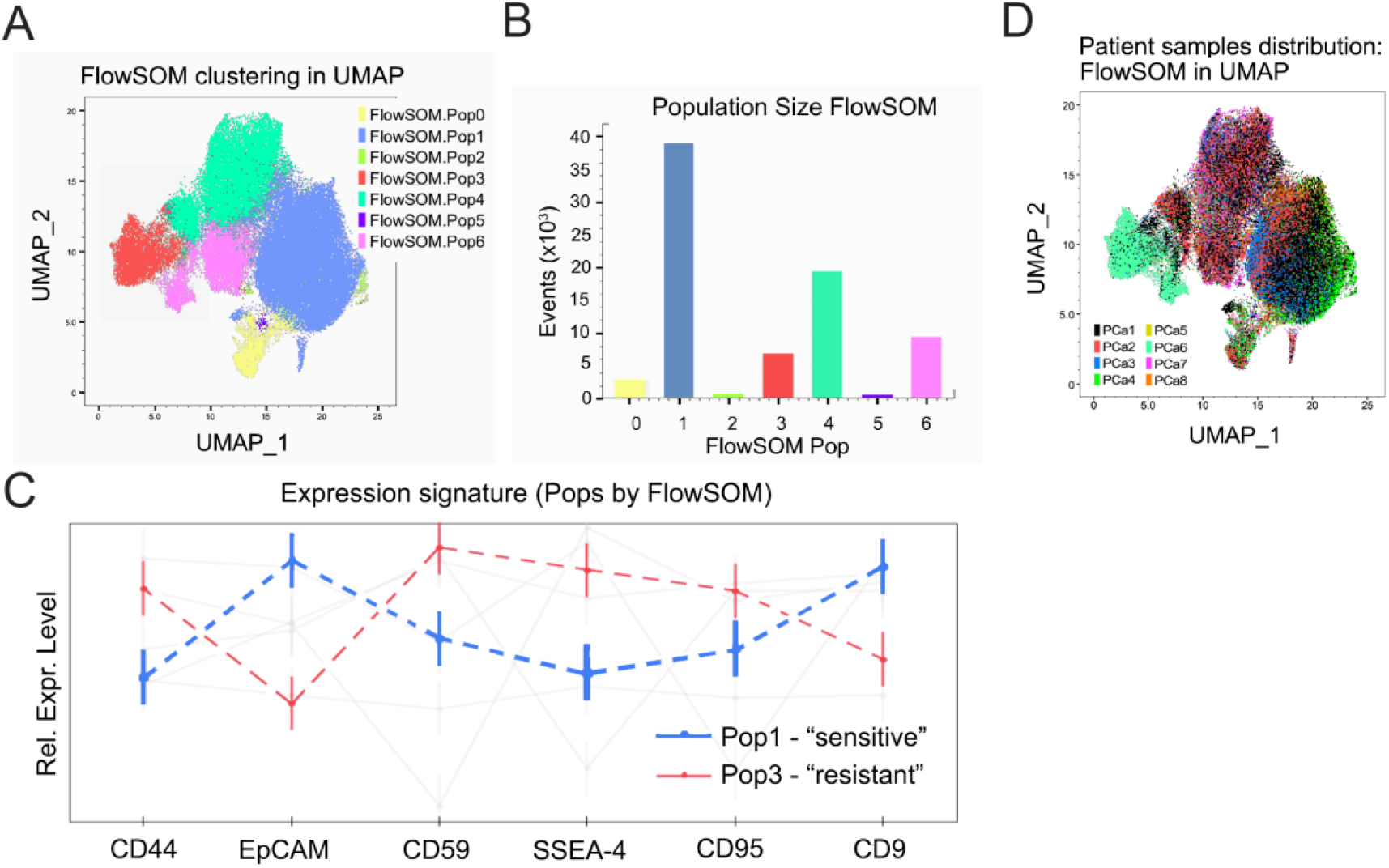
Description of the clusters and patient sample distribution within the data from dissociated primary prostate tumor samples. (A) UMAP visualization of FlowSOM clustering (related to the data in Figure 3B and 3C). (B) The bar plot shows the number of cells within each FlowSOM population. (C) The lines indicate the expression profile of the six most deregulated antigens in “sensitive” population 1 (blue) and “resistant” population 3 (red) determined by FlowSOM in prostate cancer patient samples. (D) UMAP visualization of the distribution of patient samples within the whole map determined by FlowSOM. Each color denotes a different patient sample.

**Figure S3.**
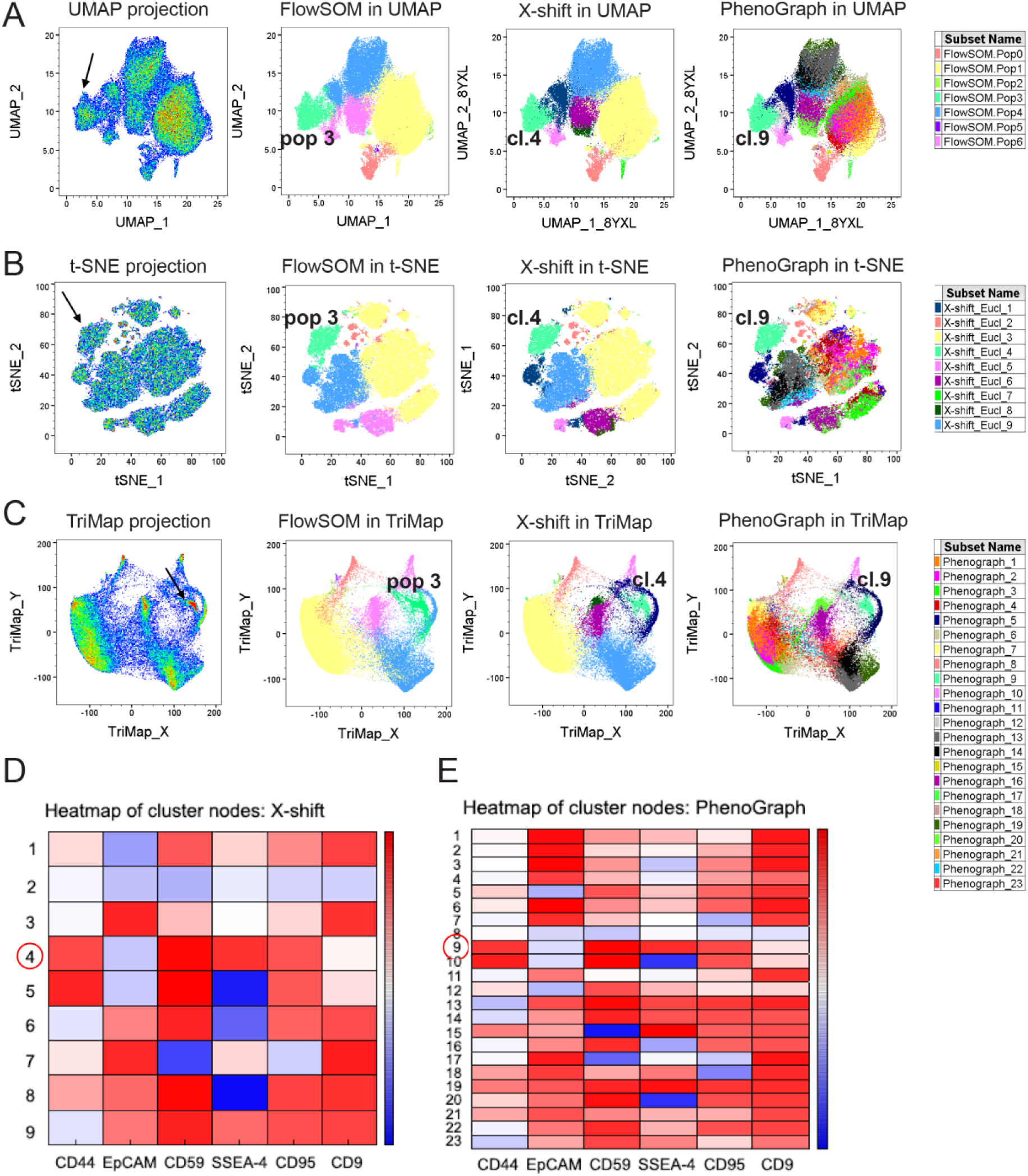
Multiple clustering and visualization analysis of the data from dissociated primary prostate tumor samples. The plots represent the clustering and visualization of clusters using different algorithms. UMAP (A), tSNE (B) and TriMap (C) algorithms were utilized for dimension reduction and map creation. To verify the presence of “resistant” population/cluster three different clustering algorithms were applied: FlowSOM (related to the data in Figure 3B and 3C), X-shift (corresponding heatmap in D) and PhenoGraph (corresponding heatmap in E). All combinations of visualization and clustering algorithms resulted in the identification of the desired population manifesting the same expression profile based on the six most deregulated surface antigens (FlowSOM - pop 3; X-shift - cluster 4 and PhenoGraph - cluster 9).

**Figure S4.**
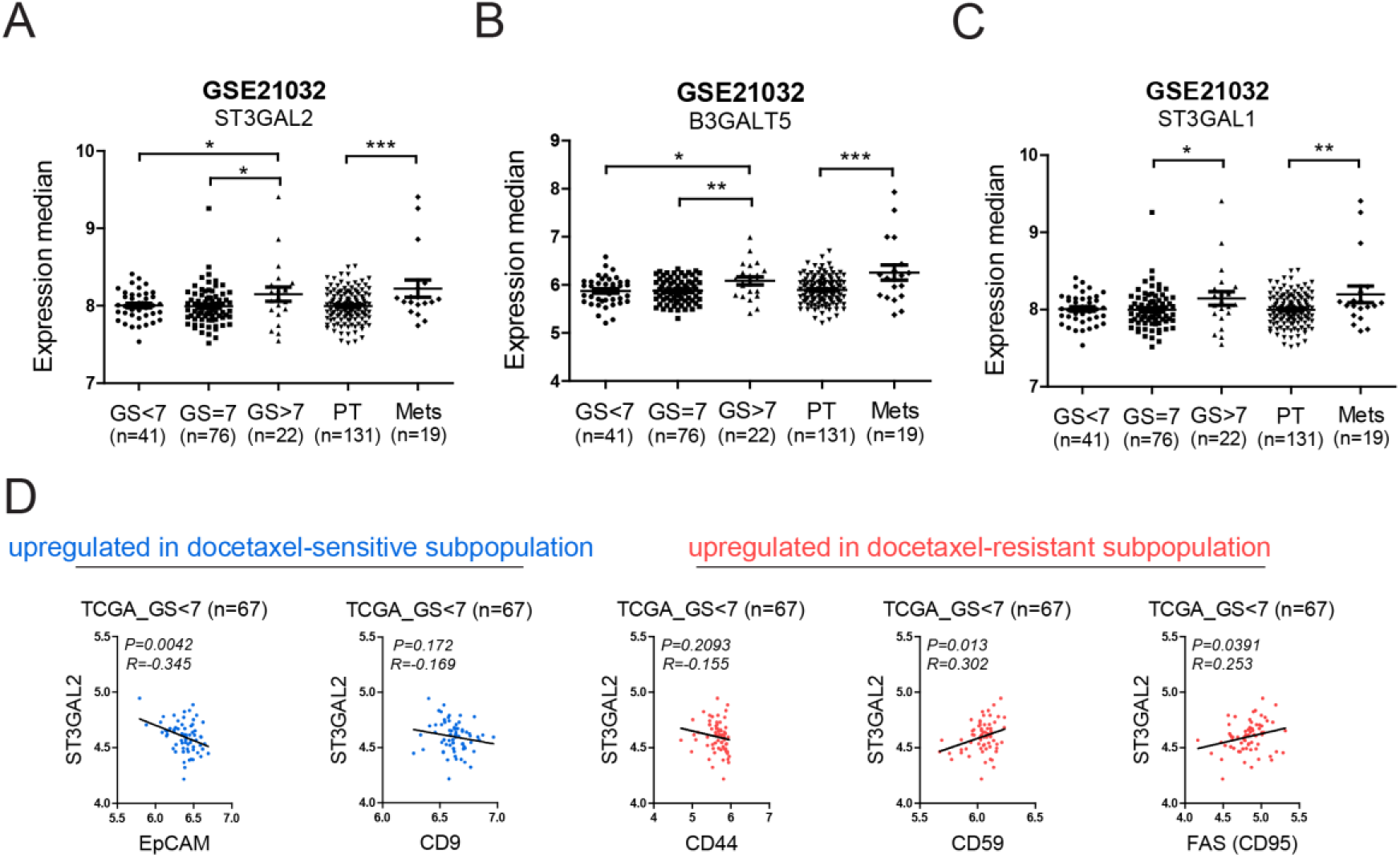
Correlation of SSEA-4 expression and disease prognosis. The expression of ST3GAL2 (A), B3GALT5 (B) and ST3GAL1(C) in the prostate cancer patient samples from various disease stage (based on the Gleason score and primary tumor (PT) or metastatic disease (Mets)) (GSE21032). (D) Correlation analysis between ST3GAL2 and surface antigens decreased (in blue) or increased (in red) in the “resistant” population of patient samples. Data portray the expression of antigens in the patients suffering from low-grade (GS<7) prostate cancer disease (n=67) (TCGA).

## Supplementary Tables

**Table S1.** LegendScreen - % of positivity.

|  | **MARKER** | **DU145** | **DU145 DOC** | **PC3** | **PC3 DOC** |
| --- | --- | --- | --- | --- | --- |
| 1 | CD1a | 1,6 | 1,8 | 1 | 2,2 |
| 2 | CD1b | 1,1 | 1,6 | 0,2 | 1 |
| 3 | CD1c | 1,1 | 0,9 | 0,1 | 1 |
| 4 | CD1d | 0,7 | 0,7 | 0 | 1,9 |
| 5 | CD2 | 1,1 | 0,9 | 0,1 | 0,9 |
| 6 | CD3 | 0,7 | 0,7 | 0,3 | 1,4 |
| 7 | CD4 | 1,3 | 1,2 | 0,2 | 0,7 |
| 8 | CD5 | 1,2 | 1,9 | 0,2 | 1,1 |
| 9 | CD6 | 1,5 | 1,9 | 0,3 | 1,3 |
| 10 | CD7 | 1,6 | 0,7 | 0,3 | 1,9 |
| 11 | CD8a | 1,7 | 1,9 | 0,1 | 1,3 |
| 12 | CD9 | 87,8 | 97,9 | 67,4 | 92,3 |
| 13 | CD10 | 1,5 | 1,9 | 0,8 | 3,2 |
| 14 | CD11a | 0,9 | 0,8 | 0,1 | 0,7 |
| 15 | CD11b | 0,9 | 0,8 | 0,1 | 1,1 |
| 16 | CD11b | 1,3 | 0,9 | 0,2 | 1 |
| 17 | CD11c | 1,4 | 1,4 | 0,2 | 0,9 |
| 18 | CD13 | 2,4 | 2,6 | 36,7 | 82 |
| 19 | CD14 | 0,6 | 0,8 | 0,7 | 2,8 |
| 20 | CD15 | 34 | 2,3 | 0,2 | 1,9 |
| 21 | CD16 | 2,2 | 1,9 | 0,1 | 1,4 |
| 22 | CD18 | 2,8 | 2,2 | 0,3 | 2 |
| 23 | CD19 | 3,6 | 4,5 | 0,6 | 2,6 |
| 24 | CD20 | 0,7 | 0,6 | 0,2 | 1,5 |
| 25 | CD21 | 1,5 | 0,8 | 0,4 | 21,4 |
| 26 | CD22 | 1,1 | 1 | 0,1 | 0,6 |
| 27 | CD23 | 0,9 | 0,7 | 0,2 | 0,5 |
| 28 | CD24 | 31,3 | 15,5 | 39,6 | 74,4 |
| 29 | CD25 | 1,2 | 1,1 | 0,3 | 1,1 |
| 30 | CD26 | 1,2 | 1 | 2,7 | 57,2 |
| 31 | CD27 | 1,1 | 1 | 0,3 | 1,1 |
| 32 | CD28 | 1,5 | 1,5 | 0,1 | 0,7 |
| 33 | CD29 | 99,2 | 99,2 | 98,5 | 99,1 |
| 34 | CD30 | 2,8 | 2 | 0,7 | 2 |
| 35 | CD31 | 3 | 3,8 | 0,3 | 2,6 |
| 36 | CD32 | 0,5 | 0,4 | 0,2 | 0,9 |
| 37 | CD33 | 0,7 | 0,6 | 0,1 | 1 |
| 38 | CD34 | 0,8 | 0,9 | 0,1 | 0,6 |
| 39 | CD35 | 1,2 | 0,5 | 0 | 0,8 |
| 40 | CD36 | 1,1 | 0,5 | 0,2 | 2,4 |
| 41 | CD38 | 15,4 | 6,2 | 0,1 | 1 |
| 42 | CD39 | 1,7 | 1,4 | 0,2 | 1,6 |
| 43 | CD40 | 3,8 | 2,4 | 0,5 | 6,6 |
| 44 | CD41 | 3,3 | 2,4 | 0,2 | 2,4 |
| 45 | CD42b | 4,1 | 3,7 | 0,2 | 1,6 |
| 46 | CD43 | 48,8 | 18 | 0,6 | 4,6 |
| 47 | CD44 | 66,6 | 99,3 | 97,3 | 99,9 |
| 48 | CD45 | 0,9 | 1 | 0,1 | 0,7 |
| 49 | CD45RA | 0,5 | 0,3 | 0,1 | 1,1 |
| 50 | CD45RB | 0,3 | 0,4 | 0,1 | 0,8 |
| 51 | CD45RO | 0,5 | 0,6 | 0,1 | 1,2 |
| 52 | CD46 | 63,9 | 13,6 | 0,9 | 12,6 |
| 53 | CD47 | 98,4 | 96,9 | 96,6 | 97,3 |
| 54 | CD48 | 5,3 | 3,1 | 0,7 | 2 |
| 55 | CD49a | 10,7 | 6,8 | 0,7 | 2,5 |
| 56 | CD49c | 99,5 | 99,7 | 95 | 99,3 |
| 57 | CD49d | 48,8 | 31,3 | 3,6 | 21,9 |
| 58 | CD49e | 96,2 | 75,5 | 19,5 | 44,1 |
| 59 | CD49f | 84,4 | 51 | 87,6 | 95,3 |
| 60 | CD50 | 1 | 0,9 | 0,2 | 0,8 |
| 61 | CD51 | 55,8 | 32,6 | 13,8 | 13,4 |
| 62 | CD51,CD61 | 1,4 | 1,4 | 0,5 | 1,6 |
| 63 | CD52 | 0,7 | 0,9 | 0,3 | 2,4 |
| 64 | CD53 | 0,8 | 1,3 | 0,3 | 0,8 |
| 65 | CD54 | 99,8 | 99,7 | 41,6 | 89,5 |
| 66 | CD55 | 99,9 | 96,7 | 98,9 | 100 |
| 67 | CD56 | 41 | 1,7 | 2,4 | 18,6 |
| 68 | CD57 | 2 | 0,9 | 1,9 | 31,5 |
| 69 | CD58 | 98,6 | 97,7 | 91 | 98,6 |
| 70 | CD59 | 98,4 | 97,4 | 97 | 96,6 |
| 71 | CD61 | 2,5 | 2,6 | 0,9 | 2,9 |
| 72 | CD62E | 0,8 | 0,9 | 0,1 | 0,9 |
| 73 | CD62L | 0,9 | 0,9 | 0,1 | 0,9 |
| 74 | CD62P | 0,9 | 0,9 | 0,2 | 0,5 |
| 75 | CD63 | 67,4 | 92,6 | 84,7 | 95,6 |
| 76 | CD64 | 1,4 | 1,4 | 0,4 | 1,6 |
| 77 | CD66a/c/e | 0,8 | 0,8 | 5,6 | 9,1 |
| 78 | CD66b | 0,7 | 0,4 | 0,5 | 0,9 |
| 79 | CD69 | 0,8 | 1,1 | 0,6 | 1,3 |
| 80 | CD70 | 67,4 | 98 | 53,5 | 91,6 |
| 81 | CD71 | 96,6 | 96,4 | 99,3 | 99,9 |
| 82 | CD73 | 96,2 | 90,5 | 26,4 | 70,4 |
| 83 | CD74 | 8,2 | 4,7 | 1,4 | 10,1 |
| 84 | CD79b | 1,1 | 1,6 | 0,3 | 1,7 |
| 85 | CD80 | 0,9 | 1,2 | 0,1 | 1,2 |
| 86 | CD81 | 96,6 | 98,1 | 90,9 | 97,4 |
| 87 | CD82 | 6,3 | 6,7 | 1,8 | 10,1 |
| 88 | CD83 | 1,4 | 2,1 | 0,5 | 2 |
| 89 | CD84 | 0,9 | 0,8 | 0,5 | 1,3 |
| 90 | CD85 | 0,6 | 0,6 | 0,2 | 0,6 |
| 91 | CD85d | 1 | 1,1 | 0,4 | 0,9 |
| 92 | CD85 | 1,7 | 1,8 | 2,9 | 2,1 |
| 93 | CD85h | 1,3 | 1,6 | 1,5 | 9,8 |
| 94 | CD85 | 0,7 | 1,2 | 1,6 | 2 |
| 95 | CD85k | 1,2 | 1,2 | 0,4 | 1,1 |
| 96 | CD86 | 0,9 | 1,1 | 0,1 | 1,1 |
| 97 | CD87 | 2,8 | 3 | 0,4 | 2,2 |
| 98 | CD88 | 1,3 | 1,5 | 0,2 | 1,6 |
| 99 | CD89 | 2,6 | 3,1 | 0,2 | 2,5 |
| 100 | CD90 | 3 | 2,8 | 0,3 | 2,8 |
| 101 | CD93 | 5,1 | 5 | 0,5 | 3,8 |
| 102 | CD94 | 3,6 | 3,4 | 0,2 | 2,3 |
| 103 | CD95 | 34,4 | 88,2 | 1 | 43 |
| 104 | CD96 | 2 | 2,9 | 0,1 | 1,5 |
| 105 | CD97 | 78,7 | 84,6 | 50,2 | 95,7 |
| 106 | CD99 | 98,8 | 98,9 | 98,5 | 99,1 |
| 107 | CD100 | 1,9 | 2 | 0,3 | 1,3 |
| 108 | CD101 | 1,5 | 2 | 0,3 | 0,6 |
| 109 | CD102 | 1,2 | 1,3 | 0,1 | 2,1 |
| 110 | CD103 | 2 | 1,8 | 0,3 | 1,4 |
| 111 | CD104 | 2,1 | 1,6 | 31,6 | 37,2 |
| 112 | CD105 | 2,6 | 2,9 | 0,4 | 2,1 |
| 113 | CD106 | 3,5 | 2,4 | 0,2 | 1,8 |
| 114 | CD107a | 11,8 | 20,2 | 2 | 21 |
| 115 | CD108 | 5,5 | 2,3 | 7,9 | 43,2 |
| 116 | CD109 | 1,8 | 2,3 | 39,5 | 76,4 |
| 117 | CD111 | 12,8 | 42,6 | 0,5 | 20,2 |
| 118 | CD112 | 94,9 | 93,7 | 20,3 | 93,5 |
| 119 | CD114 | 1,4 | 1,3 | 0,1 | 0,9 |
| 120 | CD115 | 1 | 0,8 | 0,4 | 0,7 |
| 121 | CD116 | 2,6 | 2,5 | 0,3 | 1,6 |
| 122 | CD117 | 1,6 | 1,5 | 0,1 | 0,8 |
| 123 | CD119 | 4,6 | 3,3 | 0,5 | 4 |
| 124 | CD122 | 2 | 1,4 | 0,2 | 1,2 |
| 125 | CD123 | 1,9 | 1,7 | 0,4 | 1,2 |
| 126 | CD124 | 1,6 | 1,3 | 0,8 | 3,3 |
| 127 | CD126 | 1,9 | 1,2 | 0,1 | 1,6 |
| 128 | CD127 | 2 | 2,1 | 0,4 | 9,3 |
| 129 | CD129 | 1,1 | 1,5 | 0,6 | 2,7 |
| 130 | CD131 | 1,9 | 2,1 | 0,1 | 1,3 |
| 131 | CD132 | 0,5 | 0,8 | 0,4 | 0,7 |
| 132 | CD134 | 1,6 | 1,9 | 0,2 | 0,9 |
| 133 | CD135 | 1,9 | 1,7 | 0,1 | 1 |
| 134 | CD137 | 1,8 | 1,8 | 0,1 | 1,3 |
| 135 | 4-1BB Ligand | 3,2 | 3,3 | 0,3 | 1,7 |
| 136 | CD138 | 5,3 | 3,7 | 0,5 | 7,8 |
| 137 | CD140a | 2,8 | 2,3 | 0,2 | 2,1 |
| 138 | CD140b | 2,4 | 2,5 | 0,3 | 2,4 |
| 139 | CD141 | 8,6 | 2,8 | 3,3 | 2,6 |
| 140 | CD143 | 2,8 | 2,1 | 0,5 | 4 |
| 141 | CD144 | 1,1 | 1,4 | 0,4 | 2,9 |
| 142 | CD146 | 91,4 | 93,4 | 34,4 | 87,8 |
| 143 | CD148 | 3 | 1,8 | 0,4 | 1,3 |
| 144 | CD150 | 1,3 | 1,4 | 0,1 | 0,9 |
| 145 | CD152 | 1,2 | 1 | 0,2 | 0,9 |
| 146 | CD154 | 1,9 | 1,3 | 0,3 | 1 |
| 147 | CD155 | 100 | 99,8 | 88 | 99,4 |
| 148 | CD156c | 98,4 | 97,2 | 86,8 | 98,4 |
| 149 | CD158 | 1,9 | 1,5 | 0,5 | 4,4 |
| 150 | CD158b | 1,2 | 1 | 0,3 | 2,3 |
| 151 | CD158d | 4,6 | 3,5 | 0,4 | 3,1 |
| 152 | CD158e1 | 1,8 | 1,6 | 0,1 | 1,3 |
| 153 | CD158f | 2,2 | 2,3 | 0,3 | 1,9 |
| 154 | CD161 | 2,9 | 2,7 | 0,1 | 1,2 |
| 155 | CD162 | 1,7 | 1,5 | 0,2 | 0,7 |
| 156 | CD163 | 2,3 | 1,7 | 0,4 | 1,9 |
| 157 | CD164 | 65,5 | 63,3 | 60 | 97,9 |
| 158 | CD165 | 80 | 43,4 | 13,4 | 84,3 |
| 159 | CD166 | 73,7 | 99,8 | 98,4 | 99,8 |
| 160 | CD167a | 7,4 | 3,9 | 2,2 | 2,5 |
| 161 | CD169 | 4,2 | 2,3 | 0,7 | 2,3 |
| 162 | CD170 | 34,2 | 16,3 | 63,6 | 89,8 |
| 163 | CD172a | 11,9 | 13,6 | 18,3 | 85,6 |
| 164 | CD172b | 1,2 | 1 | 0,3 | 1,8 |
| 165 | CD172g | 2 | 2 | 0,4 | 1,6 |
| 166 | CD178 | 2,6 | 1,9 | 0,3 | 2 |
| 167 | CD179a | 1,7 | 1,9 | 0,3 | 1,1 |
| 168 | CD179b | 1,2 | 1,2 | 0,2 | 0,9 |
| 169 | CD180 | 1,2 | 1,1 | 0,1 | 1,1 |
| 170 | CD181 | 1 | 1,2 | 0,4 | 1,4 |
| 171 | CD182 | 1,3 | 1,5 | 0,4 | 1,8 |
| 172 | CD183 | 3,4 | 2,5 | 0,4 | 3,4 |
| 173 | CD184 | 1,8 | 1,5 | 0,6 | 4,1 |
| 174 | CD193 | 0,9 | 1 | 0,5 | 2,1 |
| 175 | CD195 | 5,1 | 5,8 | 3,4 | 5,7 |
| 176 | CD196 | 1,1 | 1,3 | 0,5 | 3,7 |
| 177 | CD197 | 0,9 | 1,7 | 0,8 | 1,8 |
| 178 | CD200 | 1,5 | 1,2 | 0,4 | 0,8 |
| 179 | CD200R | 5,4 | 4,8 | 0,7 | 2,8 |
| 180 | CD201 | 13,7 | 64,3 | 23,3 | 86,2 |
| 181 | CD202b | 1,6 | 1,8 | 0,5 | 1,7 |
| 182 | CD203c | 1,4 | 1,7 | 0,7 | 2,1 |
| 183 | CD205 | 1,2 | 1,7 | 0,3 | 1,6 |
| 184 | CD206 | 1,5 | 1,5 | 0,5 | 1,2 |
| 185 | CD207 | 1,4 | 1,5 | 0,2 | 1,4 |
| 186 | CD209 | 0,7 | 0,8 | 0,5 | 2,1 |
| 187 | CD210 | 1 | 1 | 0,8 | 1,6 |
| 188 | CD213α2 | 1,2 | 2,2 | 1,1 | 56,3 |
| 189 | CD215 | 1 | 1,7 | 0,5 | 14,1 |
| 190 | CD218a | 2 | 1,6 | 0,4 | 4,3 |
| 191 | CD220 | 3,3 | 1 | 1,9 | 35,5 |
| 192 | CD221 | 17,3 | 6,7 | 0,5 | 5 |
| 193 | CD226 | 6,3 | 4,1 | 0,3 | 3,5 |
| 194 | CD229 | 6 | 4,4 | 0,3 | 3,9 |
| 195 | CD231 | 3,2 | 3,6 | 0,5 | 3 |
| 196 | CD235ab | 3,6 | 3,3 | 1,7 | 4,8 |
| 197 | CD243 | 2,7 | 54,5 | 0,7 | 60,5 |
| 198 | CD244 | 3,9 | 3,6 | 0,1 | 3 |
| 199 | CD245 | 11,7 | 10,4 | 3,6 | 20,2 |
| 200 | CD252 | 3,1 | 3,7 | 0,3 | 9 |
| 201 | CD253 | 3,1 | 3,7 | 0,2 | 1,8 |
| 202 | CD254 | 4,3 | 5,5 | 0,6 | 4,3 |
| 203 | CD255 | 1,5 | 2,1 | 0,3 | 1,2 |
| 204 | CD257 | 12,1 | 3,7 | 1 | 24,4 |
| 205 | CD258 | 1,5 | 1,3 | 0,2 | 2,1 |
| 206 | CD261 | 9,2 | 5,4 | 0,5 | 6,1 |
| 207 | CD262 | 89,3 | 44,8 | 6,7 | 39,7 |
| 208 | CD263 | 4,2 | 4,6 | 0,3 | 3,3 |
| 209 | CD266 | 41,4 | 37,2 | 1,3 | 27,8 |
| 210 | CD267 | 1,7 | 1,8 | 0,9 | 1,6 |
| 211 | CD268 | 4,5 | 3,4 | 0,4 | 3,8 |
| 212 | HVEM | 3,4 | 3,2 | 0,3 | 3,6 |
| 213 | CD271 | 6,1 | 6 | 0,5 | 5,4 |
| 214 | CD273 | 0,8 | 1,1 | 0,3 | 1,3 |
| 215 | CD274 | 3,4 | 11,1 | 14,9 | 20,1 |
| 216 | CD275 | 4,4 | 3,1 | 0,7 | 17,7 |
| 217 | CD276 | 89,7 | 96 | 69,8 | 95 |
| 218 | CD277 | 10,6 | 8,6 | 0,4 | 5,3 |
| 219 | CD278 | 1,5 | 1,1 | 1 | 3,4 |
| 220 | CD279 | 5,9 | 4,6 | 0,7 | 4,9 |
| 221 | CD282 | 2,7 | 1,9 | 0,8 | 6,2 |
| 222 | CD284 | 5,1 | 5 | 2,8 | 13,4 |
| 223 | CD286 | 4,5 | 3,3 | 0,2 | 3,1 |
| 224 | CD290 | 3,6 | 3 | 0,3 | 2,4 |
| 225 | CD294 | 3,3 | 2,7 | 1,1 | 2,5 |
| 226 | CD298 | 98,5 | 97,8 | 96,2 | 97,4 |
| 227 | CD300e | 1,1 | 0,7 | 0,8 | 0,7 |
| 228 | CD300F | 2,1 | 1,9 | 0,5 | 1,8 |
| 229 | CD301 | 3,8 | 2,7 | 2,1 | 9,1 |
| 230 | CD303 | 2,6 | 1,7 | 1,1 | 5 |
| 231 | CD304 | 7,5 | 3,1 | 65,4 | 94,8 |
| 232 | CD307e | 2,7 | 1,7 | 0,9 | 5,2 |
| 233 | FcRL4 | 3,8 | 2,3 | 2,2 | 7,4 |
| 234 | CD314 | 7,2 | 5,4 | 0,7 | 5,9 |
| 235 | CD317 | 19 | 8,6 | 2,8 | 12,3 |
| 236 | CD318 | 98,9 | 95,5 | 95,7 | 99,7 |
| 237 | CD319 | 7,6 | 5,6 | 2 | 13,7 |
| 238 | CD324 | 40,3 | 2,5 | 4,9 | 15 |
| 239 | CD325 | 1,3 | 1,4 | 0,4 | 1,5 |
| 240 | CD326 | 98,7 | 67,4 | 95,7 | 98,8 |
| 241 | CD328 | 7,7 | 5,7 | 2,1 | 6,5 |
| 242 | CD334 | 10,1 | 6,6 | 0,5 | 7,4 |
| 243 | CD335 | 6,4 | 3,9 | 1 | 5,8 |
| 244 | CD336 | 8,2 | 6,2 | 0,9 | 5,5 |
| 245 | CD337 | 4,7 | 3,2 | 0,7 | 3,8 |
| 246 | CD338 | 8,9 | 6,2 | 1,2 | 8,3 |
| 247 | CD340 | 41,9 | 32,6 | 2,9 | 42,5 |
| 248 | CD344 | 8,1 | 4,5 | 0,6 | 13,4 |
| 249 | CD351 | 13,3 | 8,5 | 0,6 | 8,1 |
| 250 | CD352 | 0,9 | 1,4 | 0,1 | 1 |
| 251 | CD354 | 1,1 | 0,9 | 0,1 | 1 |
| 252 | CD355 | 1,2 | 1 | 1,1 | 2,2 |
| 253 | CD357 | 6,5 | 8,4 | 1,2 | 5,4 |
| 254 | CD360 | 14,3 | 9,2 | 2,4 | 10,3 |
| 255 | β2-microglobulin | 99 | 98,4 | 97,9 | 98,9 |
| 256 | CD272 | 2,6 | 1,7 | 2,7 | 7 |
| 257 | C3aR | 3,1 | 2,8 | 2,1 | 10,1 |
| 258 | C5L2 | 2,6 | 2,2 | 3,5 | 7,6 |
| 259 | CCR10 | 2,4 | 1,4 | 2,1 | 6,7 |
| 260 | CLEC12A | 3 | 2,6 | 1,8 | 7 |
| 261 | CLEC9A | 3 | 2,6 | 1,6 | 5,4 |
| 262 | CX3CR1 | 0,8 | 0,9 | 1,9 | 1,4 |
| 263 | CXCR7 | 2,4 | 4,1 | 1,5 | 4,1 |
| 264 | Delta Opioid Receptor | 1,6 | 1,9 | 1,3 | 3 |
| 265 | DLL1 | 11 | 5,5 | 1,6 | 7,7 |
| 266 | DLL4 | 13,4 | 7,6 | 1,6 | 9,5 |
| 267 | DR3 | 8,9 | 6,2 | 2,9 | 8,2 |
| 268 | EGFR | 97,6 | 94,3 | 52,6 | 94,2 |
| 269 | erbB3 | 16,7 | 9,2 | 6,7 | 20,6 |
| 270 | FcεRIα | 1,7 | 1,5 | 4,5 | 7,1 |
| 271 | FcRL6 | 2,2 | 2 | 3,3 | 6 |
| 272 | Galectin-9 | 3,9 | 3,3 | 0,6 | 2,5 |
| 273 | GARP | 2,4 | 1,7 | 2,4 | 5,3 |
| 274 | HLA-A,B,C | 92,3 | 94 | 85,4 | 95,4 |
| 275 | HLA-A2 | 0,7 | 0,6 | 1,5 | 1,2 |
| 276 | HLA-DQ | 2,7 | 1,6 | 1,1 | 2,6 |
| 277 | HLA-DR | 3 | 2,1 | 3,9 | 9,3 |
| 278 | HLA-E | 13,8 | 8 | 5,3 | 11,9 |
| 279 | HLA-G | 1,7 | 1,4 | 5,3 | 6,5 |
| 280 | IFN-γ R b chain | 1,7 | 1,8 | 1,8 | 4,9 |
| 281 | Ig light chain κ | 4,6 | 3,2 | 0,9 | 3,2 |
| 282 | Ig light chain λ | 1,3 | 0,9 | 0,1 | 3 |
| 283 | IgD | 1,7 | 1,8 | 0,8 | 4,5 |
| 284 | IgM | 4,5 | 3,2 | 0,9 | 3,2 |
| 285 | IL-28RA | 1,1 | 0,7 | 0,8 | 1,8 |
| 286 | Integrin α9β1 | 2,8 | 2,7 | 0,4 | 2,1 |
| 287 | integrin β5 | 48,3 | 45,9 | 2,7 | 6,1 |
| 288 | Integrin β7 | 2,1 | 1,8 | 0,9 | 0,8 |
| 289 | Jagged 2 | 4,4 | 4,2 | 0,7 | 3,2 |
| 290 | LAP | 3,8 | 3,9 | 1,2 | 2,5 |
| 291 | Lymphotoxin β Receptor | 7,8 | 10,7 | 5,5 | 40,2 |
| 292 | Mac-2 | 3,5 | 3,7 | 0,1 | 3,8 |
| 293 | MAIR-II | 4,9 | 5,5 | 0,4 | 4,5 |
| 294 | MICA/MICB | 42 | 10,2 | 1,6 | 7,1 |
| 295 | SUSD2 | 5,6 | 3,6 | 0,4 | 6,3 |
| 296 | SUSD2 | 10,2 | 6 | 0,4 | 5,1 |
| 297 | MSC | 91 | 87 | 11,4 | 76,5 |
| 298 | MSC,NPC | 11,2 | 6 | 3,7 | 50,1 |
| 299 | TNAP | 2,8 | 3,2 | 0,3 | 2,9 |
| 300 | NKp80 | 7,7 | 3,9 | 0,7 | 4,1 |
| 301 | Notch 1 | 3,7 | 3,4 | 1,6 | 4,6 |
| 302 | Notch 2 | 2,4 | 2,7 | 2,4 | 19 |
| 303 | Notch3 | 4,7 | 3,8 | 0,3 | 3,9 |
| 304 | Notch 4 | 2,6 | 2 | 0,2 | 4,9 |
| 305 | NPC | 6,7 | 5,4 | 0,3 | 4,9 |
| 306 | Podoplanin | 3,7 | 2,1 | 7,1 | 7,9 |
| 307 | Pre-BCR | 5,7 | 5,1 | 1,1 | 5 |
| 308 | PSMA | 5,1 | 5,2 | 1,1 | 4,1 |
| 309 | Siglec-10 | 3,4 | 3,8 | 0,2 | 2 |
| 310 | Siglec-8 | 2,8 | 3,3 | 0,9 | 3 |
| 311 | Siglec-9 | 3,6 | 3 | 0,6 | 2,6 |
| 312 | SSEA-1 | 10,3 | 1,1 | 1,9 | 2 |
| 313 | SSEA-3 | 1,5 | 1,6 | 0,5 | 4,1 |
| 314 | SSEA-4 | 20,4 | 73,9 | 85 | 95,9 |
| 315 | SSEA-5 | 47,2 | 48,5 | 72,2 | 89,6 |
| 316 | TCR $\gamma/\delta$ | 6,9 | 6,8 | 3,3 | 7,1 |
| 317 | TCR Vβ13.2 | 7,6 | 7,1 | 1,6 | 7,5 |
| 318 | TCR Vβ23 | 5,5 | 4,7 | 0,9 | 4,4 |
| 319 | TCR Vβ8 | 4 | 3,6 | 2,5 | 7,3 |
| 320 | TCR Vβ9 | 2,4 | 2,3 | 1,9 | 6,1 |
| 321 | TCR Vδ2 | 2,3 | 2,6 | 0,8 | 1,6 |
| 322 | Vγ9 | 3,5 | 3,7 | 0,4 | 2,5 |
| 323 | TCR Vα24-Jα18 | 2,1 | 2,1 | 0,4 | 1,8 |
| 324 | TCR Vα7.2 | 4,2 | 3,3 | 0,5 | 3,4 |
| 325 | TCR $\alpha/\beta$ | 4,1 | 3,4 | 1,8 | 3,5 |
| 326 | Tim-1 | 3 | 2,4 | 0,6 | 1,5 |
| 327 | Tim-3 | 5,4 | 4,3 | 0,4 | 2,9 |
| 328 | Tim-4 | 7,7 | 5,1 | 0,3 | 4 |
| 329 | TLT-2 | 6,5 | 5,1 | 4,5 | 5,7 |
| 330 | TRA-1-60-R | 9 | 21,8 | 5,4 | 10,9 |
| 331 | TRA-1-81 | 3,4 | 4,5 | 2,7 | 3,4 |
| 332 | TSLPR | 3,8 | 3,8 | 0,6 | 2,3 |

**Table S2.**
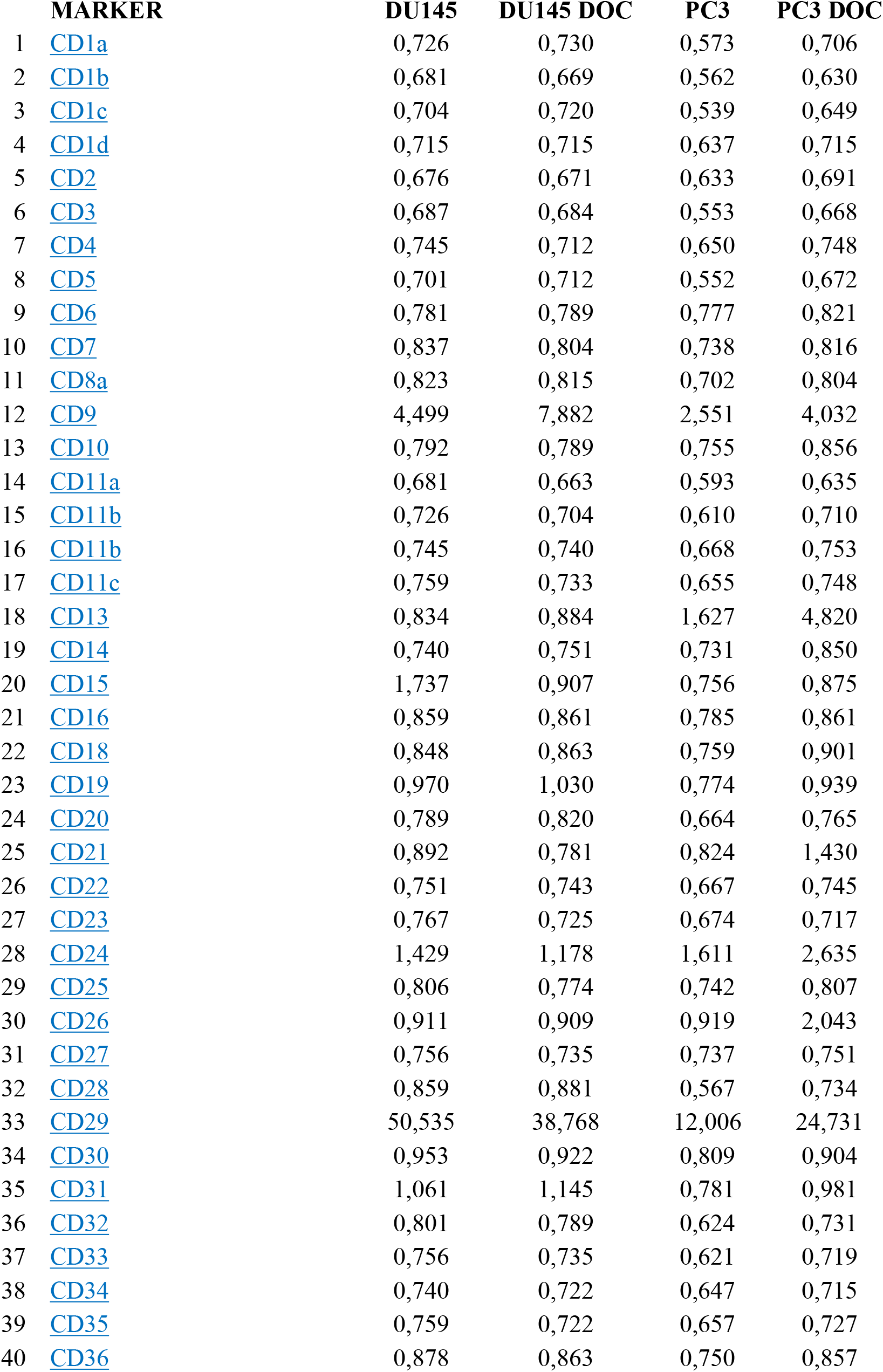
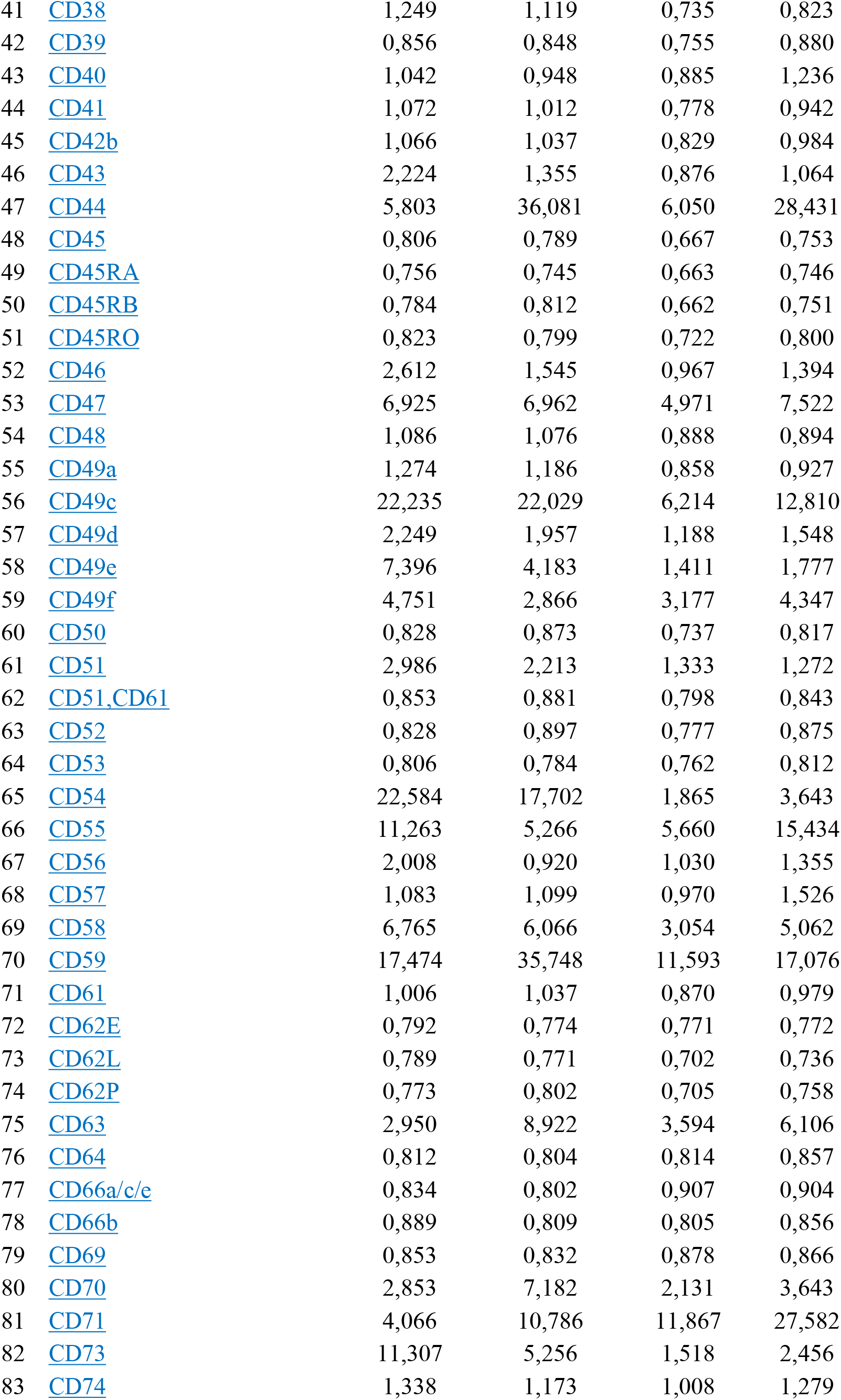
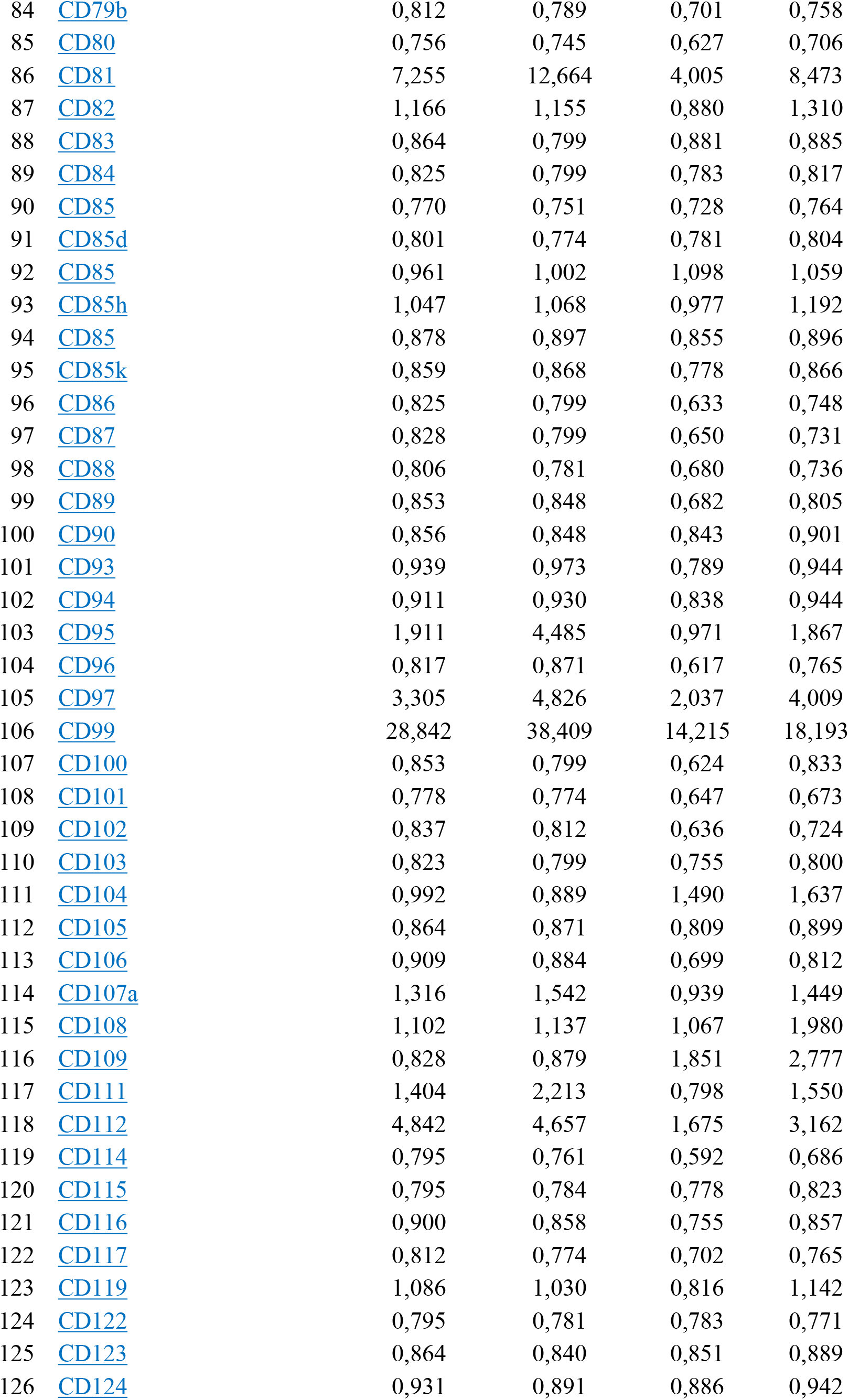
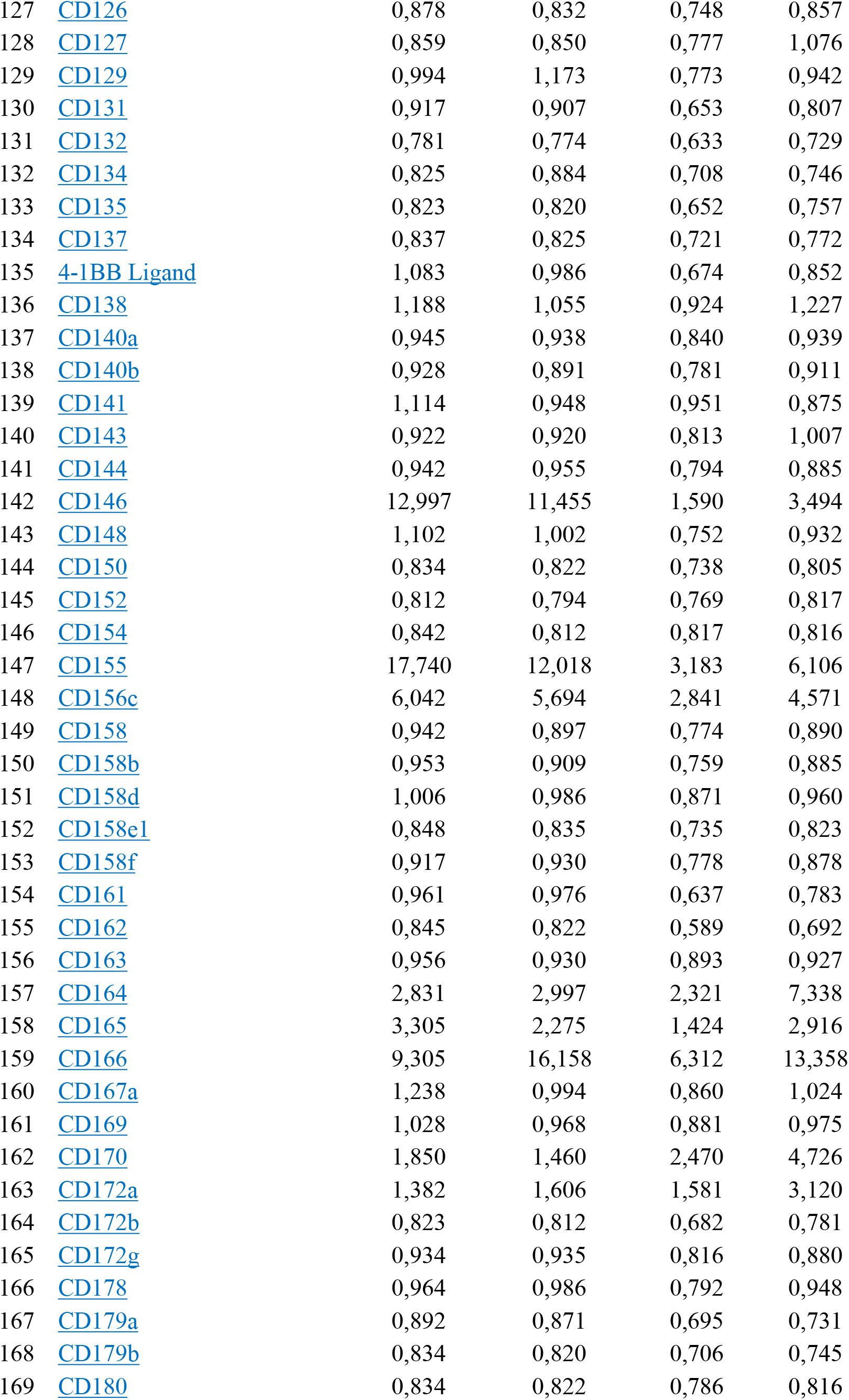
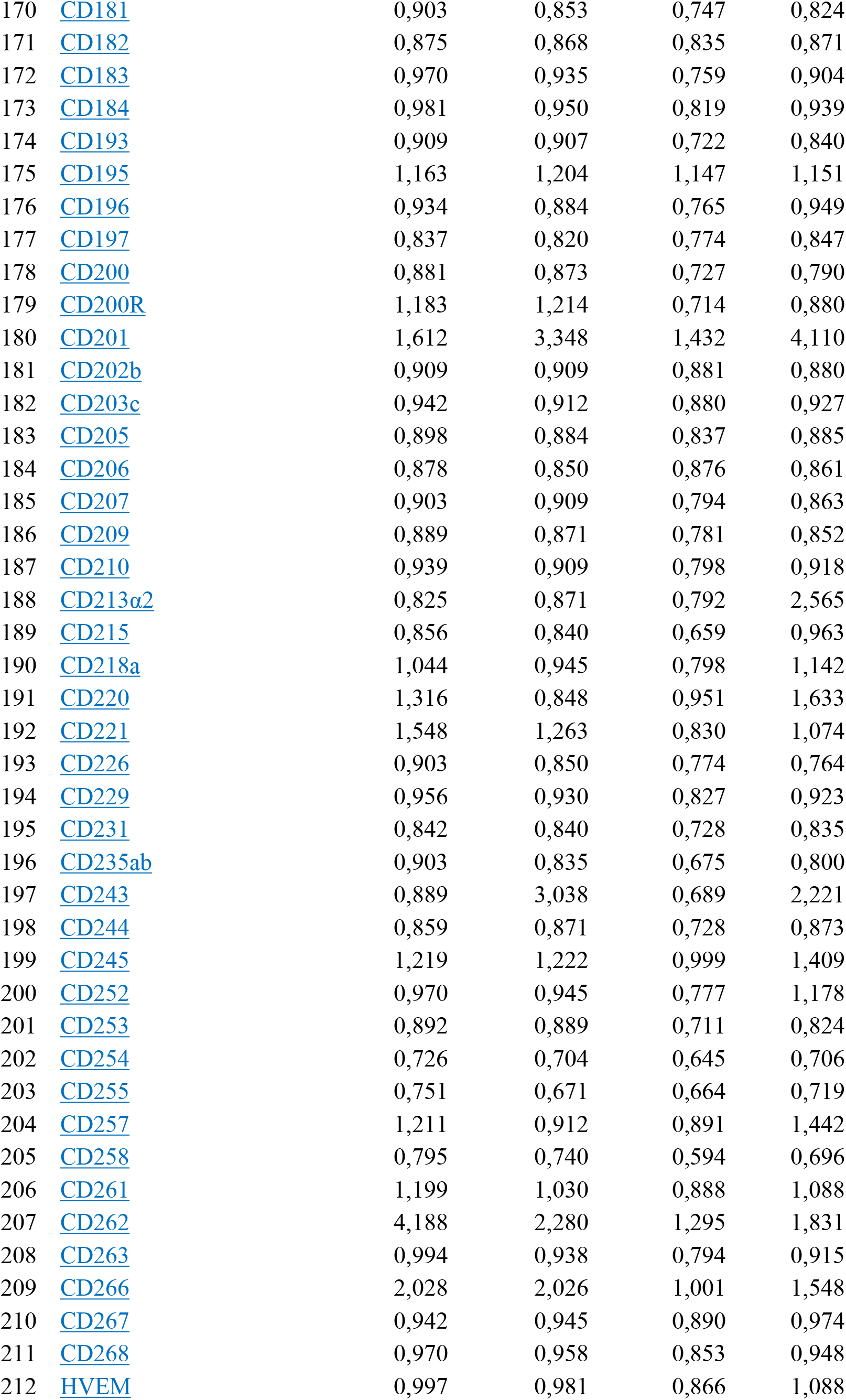
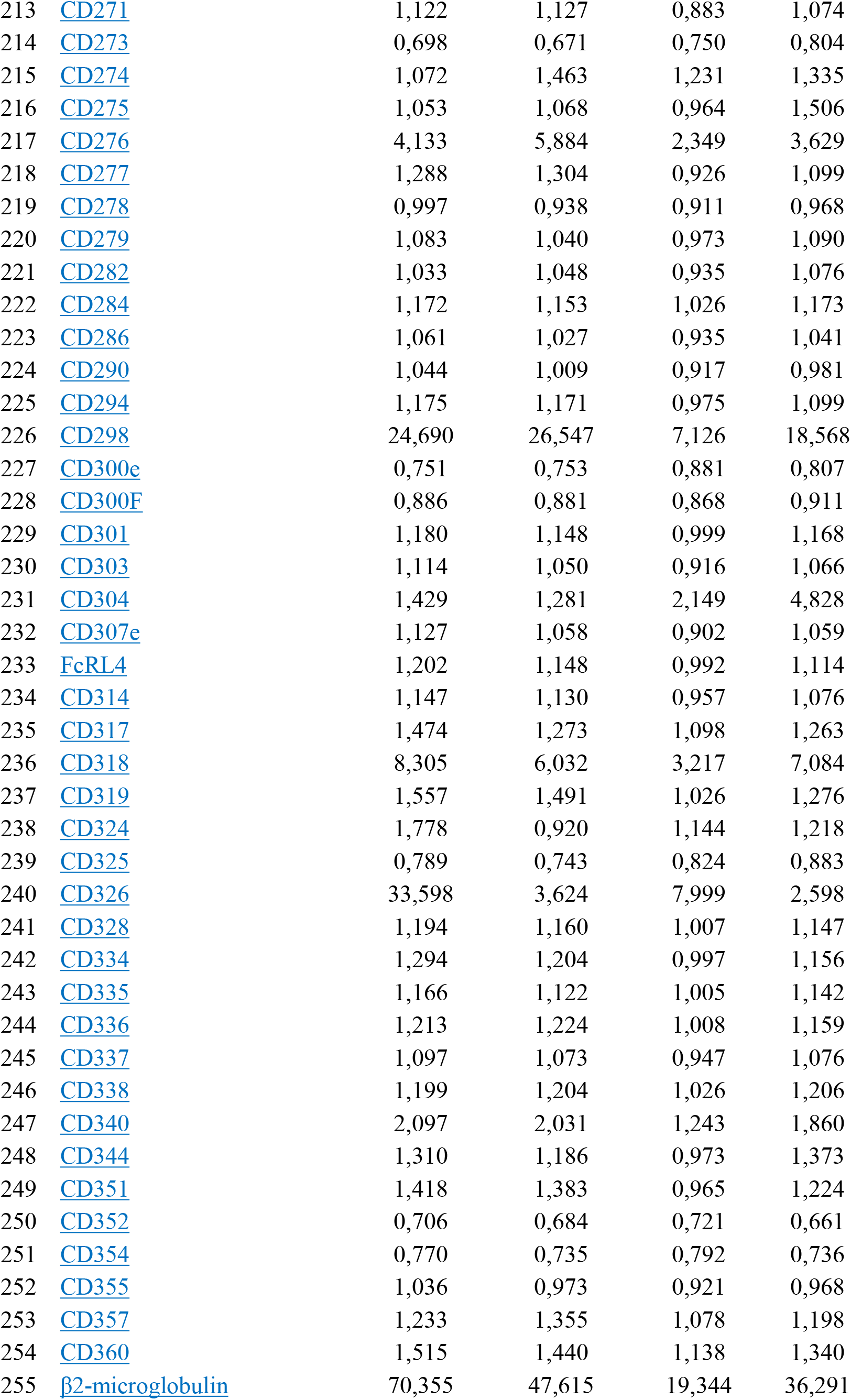
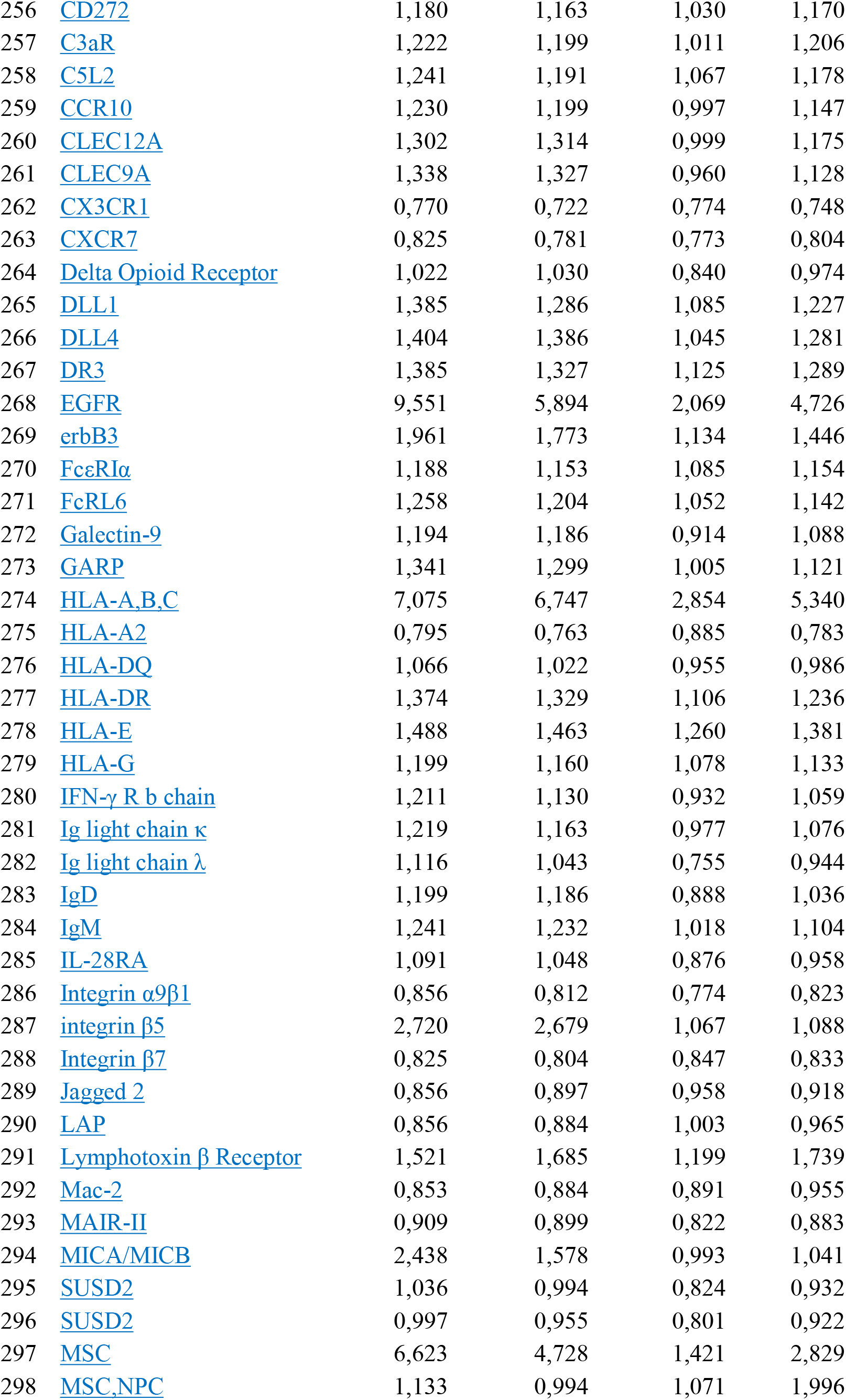
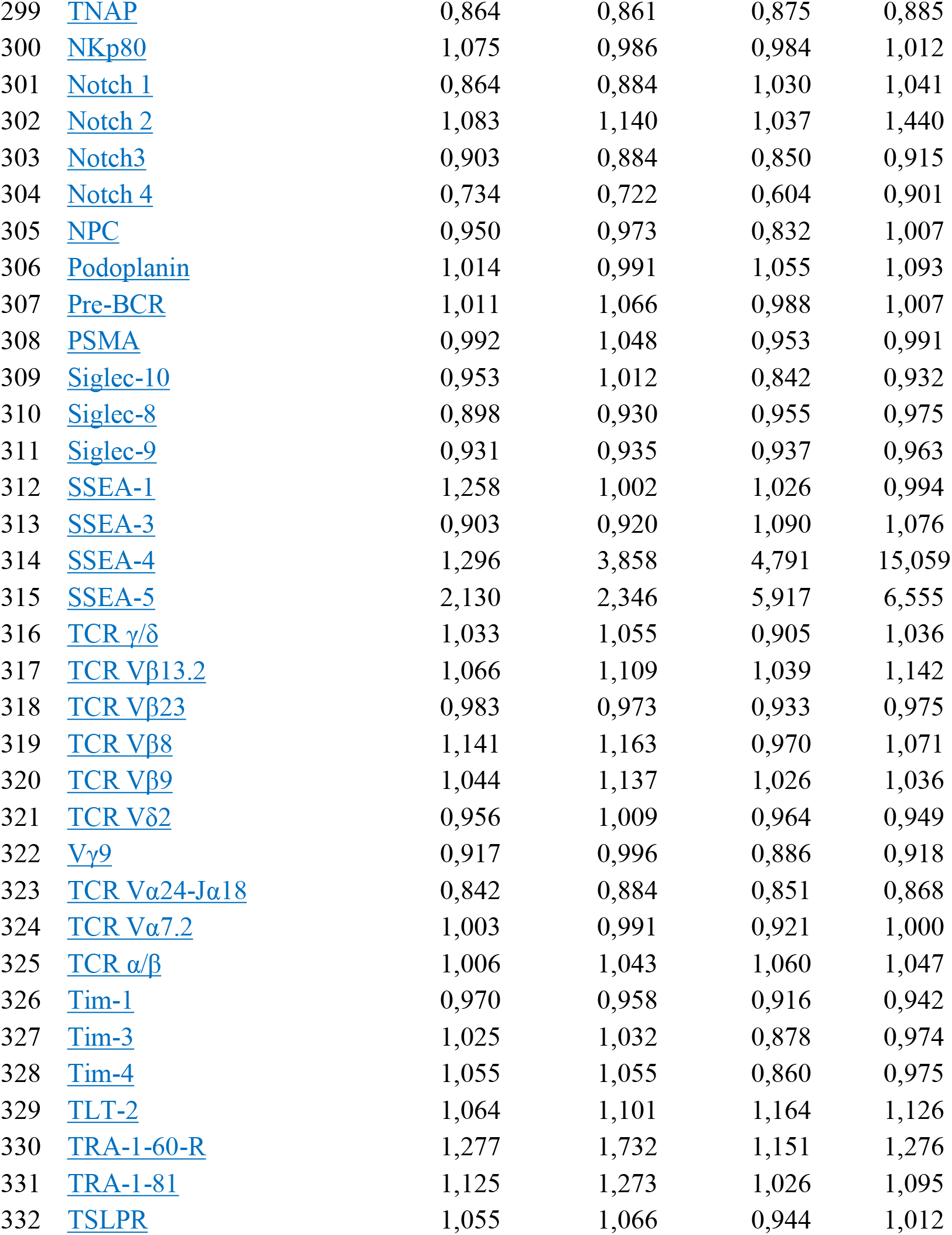
LegendScreen - MFI of 332 surface molecules (normalized to ISO).

**Table S3.** Overview of prostate cancer patient specimens.

| **Sample No.** | **Sample ID** | **Diagnosis** | **Gleason score** | **Grade group** | **TNM** |
| --- | --- | --- | --- | --- | --- |
| PCa1 | 2033 | adenocarcinoma | 4+3 | 3 | pT3a, pN0, pMX |
| PCa2 | 1981 | adenocarcinoma | 3+4 | 2 | pT3a, pN0, pMX |
| PCa3 | 2036 | adenocarcinoma | 4+3 | 3 | pT3a, pN1, pMX |
| PCa4 | 2046 | adenocarcinoma | 4+5 | 5 | pT3b, pN1, pMX |
| PCa5 | 2040 | adenocarcinoma | 4+5, GS 5 up to 5% | 5 | pT3a, pN0, pMX |
| PCa6 | 2044 | adenocarcinoma | 4+3, GS 5 up to 5% | 3 | pT3a, pN0, pMX |
| PCa7 | 2010 | adenocarcinoma | 4+3, GS 5 up to 5% | 3 | pT3a, pN1, pMX |
| PCa8 | 1591 | adenocarcinoma | 4+5 | 5 | pT3b, pN0, pMX |

| **Sample No.** | **Ki67 (%)** | **iPSA** | **Therapy** | **Surgery** |
| --- | --- | --- | --- | --- |
| PCa1 | 23 | 6,6 | without adjuvant therapy | 08.06.2020 |
| PCa2 | 2 | 4,77 | without adjuvant therapy | 13.12.2019 |
| PCa3 | 11 | 4,8 | without adjuvant therapy | 09.06.2020 |
| PCa4 | 21 | 15,44 | hormonal therapy LHRH | 23.06.2020 |
| PCa5 | 17 | 16,2 | without adjuvant therapy | 15.06.2020 |
| PCa6 | 8 | 5,32 | without adjuvant therapy | 23.06.2020 |
| PCa7 | 9 | 5,13 | without adjuvant therapy | 18.02.2020 |
| PCa8 | 4 | 9,94 | salvage RT+ LHRH | 31.01.2018 |

**Table S4.** Overview of antibodies and reagents used for (spectral) flow cytometry.

| Antigen | Conjugate | Host | Isotype and clonality | Producer | Cat.No. | Dilution |
| --- | --- | --- | --- | --- | --- | --- |
| CD9 | CF-Blue | Mouse | IgG2a,κ; clone VJ1/20 | Immunostep | 9CFB-100T | 1:20 |
| CD44 | AF532 | Rat | IgG2b; clone IM7 | eBioscience | 58-0441-82 | 1:40 |
| CD59 | PE | Mouse | IgG2a,κ; clone p282 | SONY | 2123540 | 1:40 |
| CD63 | BV650 | Mouse | IgG1,κ; clone H5C6 | Biolegend | 353026 | 1:20 |
| CD70 | BV786 | Mouse | IgG3,κ; clone Ki-24 | BD Horison | 565338 | 1:20 |
| CD71 | VioGreen | Mouse | IgG1,κ; clone REA902 | Miltenyi Biotec | 130-115-035 | 1:20 |
| CD81 | PE-Cy7 | Mouse | IgG1,κ; clone 5A6 | SONY | 2347560 | 1:40 |
| CD95 | PE-Cy5 | Mouse | IgG1,κ; clone DX2 | Biolegend | 305610 | 1:320 |
| CD97 | BV605 | Mouse | IgG1,κ; clone VIM3b | BD OptiBuild | 742447 | 1:80 |
| CD166 | PerCP-Cy5.5 | Mouse | IgG1,κ; clone 3A6 | Biolegend | 343908 | 1:80 |
| CD201 | BV711 | Rat | IgG1,κ; clone RCR-252 | BD OptiBuild | 743555 | 1:80 |
| SSEA-4 | PE-CF594 | Mouse | IgG3,κ; clone MC813-70 | BD Horison | 562487 | 1:20 |
| EpCAM | BV421 | Mouse | IgG2b,κ; clone 9C4 | Biolegend | 324220 | 1:200 |
| hCD298 | FITC | Recomb. Hu. | IgG1; clone REA217 | Miltenyi Biotec | 130-123-324 | 1:100 |
| CD31 | FITC | Mouse | IgG1,κ; clone WM59 | SONY | 2115520 | 1:100 |
| CD45 | FITC | Mouse | IgG1,κ; clone HI30 | SONY | 2120030 | 1:100 |
| CD90 | FITC | Mouse | IgG1,κ; clone Thy1 | Biolegend | 328108 | 1:100 |
| **Viability** |  |  |  |  |  |  |
| Live/Dead | Yellow |  |  | Invitrogen | L34959 | 1:500 |
|  | Green |  |  | Invitrogen | L23101 | 1:1000 |
| **Barcoding** |  |  |  |  |  |  |
| CellTrace | Violet |  |  | Invitrogen | C34557 | 1:500 -<br>1:10 000 |
|  | Far Red DDAO-SE |  |  | Invitrogen | C34553 | 1:1000 -<br>1:10 000 |
| **Others** |  |  |  |  |  |  |
| Super Bright Complete Staining Buffer |  |  |  | eBioscience | SB-4401-42 | 1:100 |

